# Unlocking the secrets of paired terminal genomic repeats utilised by the *Burkholderia* jumbo phage Matilda

**DOI:** 10.64898/2026.10.21.760668

**Authors:** Jessica M Lewis, Olivia Burns, Harry Mogg, Tanaz Mistry, Yan-Jiun Lee, Peter Weigele, Eshwar Mahenthiralingam, Yin Chen, Nichollas E Scott, Laura Baxter, Cristina S Mesquita, Richard Stark, Trevor Lithgow, Jeremy J Barr, Antonia P Sagona

## Abstract

Bacteriophages (phages) are viruses that infect bacteria. Jumbo phages have genomes exceeding 220 kb, which can support greater functional independence and complexity. This work identified vB_BceM_Matilda (φMatilda), a novel *Burkholderia*-targeting jumbo phage. Before injecting its genome into the host bacterium, φMatilda binds to the bacterial flagellum, the rotation of which is required to drive a productive phage infection. Its linear 267 kb dsDNA genome has a previously undescribed tripartite architecture, with bidirectional terminal elements of 1.3 kb and 25 kb, each flanked by distinct inverted repeats of 1.5 kb and 18 kb, respectively. These architectural elements bookend the genome. Transcriptomic and proteomic analyses tracked the phage infection process and confirmed that genes within these terminal elements are transcribed and translated during infection. The presence of bidirectional terminal sequences in related jumbo phages suggests that the tripartite genome architecture may be conserved and functionally important. This work represents the first description of this genome architecture in any organism, expanding our understanding of genome organisation and raising new questions about the replication, packaging, and evolution of jumbo phage genomes.

**Visual abstract:** 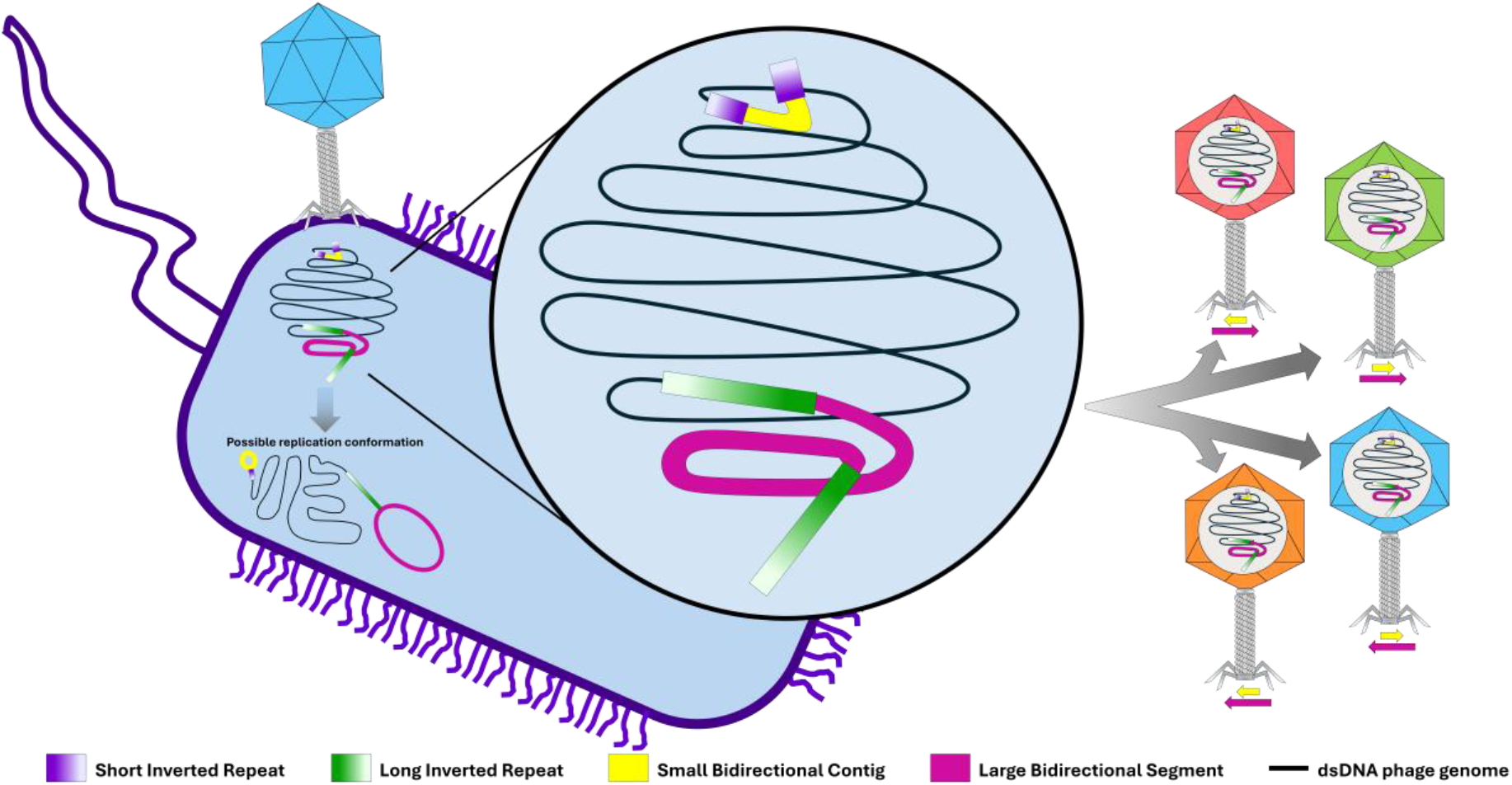

## Introduction

Viruses that infect bacteria, known as bacteriophages (phages), play important roles in modulating bacterial communities through a constant evolutionary arms race with their hosts for survival^1^. This battle has enabled phages to shape microbial communities and has increased interest in their practical applications. Phages have been used in agriculture to prevent infections of livestock and crops, improving productivity and crop yields while reducing the reliance on antibiotics. In medicine, phages have been used on a compassionate basis to treat multidrug resistant infections. Despite these current applications, much of phage diversity remains uncharacterised, limiting our understanding of phage biology^2,3^. Double stranded DNA (dsDNA) phages belonging to *Caudoviricetes* represent the majority of sequenced phages^4^ and thus serve as model systems for studying the phage infection process, including DNA replication and packaging^5–7^. Despite differences in intracellular genome conformations during infection, these tailed phages package linear DNA within their capsid^8^.

Following injection into the bacterial host cell, phage genomes form different conformations to both enable replication and evade host defence systems. For example, multiple phages convert their linear genomes into circular forms to protect exposed termini from degradation. This can occur through annealing of cohesive (*cos*) ends^9,10^ or through recombination of direct terminal repeats (DTRs) that vary in length^5,11,12^. Circularised phage genomes typically replicate via rolling circle replication and are packaged as either one full length genome (*cos* phages) or as concatemeric DNA, which can include up to 110% of the genome (DTR phages)^13^.

In contrast, certain phages employ alternative strategies to avoid degradation without circularisation, such as coliphage T4, which employs DNA base modification to resist host endonucleases^14–16^. Coliphage T7 uses a staged injection process, allowing expression of anti-defence proteins including the Ocr protein^17–19^, which shuts down the endogenous restriction/modification systems thereby protecting its genome. Phage T5 adopts a similar staged injection strategy to enable synthesis of defence inhibitors^20^ although it ultimately circularises its genome. Other phages, such as φ29, inject cargo proteins that cap the DNA termini, protecting the DNA from host cell exonucleases and facilitating replication^21,22^. While these strategies illustrate the diverse mechanisms utilised by tailed phages, these models are largely derived from model phages with relatively small genomes.

Jumbo phages have genomes exceeding 200 kb and, while they represent a small proportion of known phages, their larger genomes confer greater genetic capabilities^23^. Compared to their smaller counterparts, jumbo phages produce larger and more complex virions^24^, enabling greater independence from the host. This includes encoding their own replication machinery^6,25–27^ and complex systems that mitigate bacterial defence systems^6,23^. Despite the relatively small number of jumbo phages that have been isolated to date, they display considerable functional diversity. Notably, some jumbo phages have evolved the ability to form nucleus-like compartments within the bacterial cell that protect the phage genome from host-mediated degradation^26,28^. As such, jumbo phages are an underexplored reservoir for unique biological mechanisms.

Bacteria belonging to the *Burkholderia cepacia* complex (BCC) are widely found in soil and water, and can cause infections in plants, including crops such as onions and rice^29^, as well as opportunistic lung infections in people with cystic fibrosis (CF)^30^. Phages targeting *Burkholderia* are studied both for their ability to treat human infections and to control plant diseases^2,31,32^. However, compared with other pathogenic Gram-negative bacteria, relatively few *Burkholderia* phages have been isolated and sequenced, and only seven of these are jumbo phages^2,31,33–36^. In this study, a novel dsDNA jumbo phage targeting *Burkholderia cenocepacia* was isolated that possesses a previously undescribed genome architecture composed of large bidirectional terminal elements contained within long inverted repeats. While this is the first characterisation of such genome architecture, phylogenetic analysis of publicly available phage genomes suggests that similar architecture may be more widespread amongst other jumbo phages. This unique architecture raises questions about how the genome is packaged into the capsid, how the genome termini are protected following ejection into the host cell, how replication initiates, and whether the architecture contributes to the evasion of host defences. Genomic, transcriptomic, and proteomic analyses were undertaken to characterise this unusual architecture and to assess its biological significance. Together, these findings reveal a previously unrecognised architecture for genome organisation and demonstrate that the associated terminal elements are actively deployed during the infection cycle.

## Methods

### Organisms and culture conditions

The bacterial strains utilised in this study are listed in Supplementary Table 1^37–52^ and encompassed 31 *Burkholderia* strains representative of 15 species. All species were routinely cultured in lysogeny broth (LB; Invitrogen, USA). Liquid cultures were incubated at 37°C, 180 rpm for 16 hours. For solid media, LB was supplemented with 1% agar (w/v, VWR Chemicals, USA) and strains were grown for 24-48 hours at 37°C. Phages were propagated at 30°C for 20 hours unless otherwise specified. Plaque assays were performed using 100 µL of overnight bacterial culture and 100 µL of phage sample combined with 3 mL of 0.5% (w/v) LB agar + 2mM MgSO_4_ (Sigma, Japan) + 2mM CaCl_2_ (Sigma, Japan) and poured over a 1% LB agar base. Strains harbouring pSCrhaB2 were cultured with 100 µg/mL trimethoprim (APExBIO, USA), and expression was induced with 0.1% L-Rhamnose (w/v, Sigma, Japan).

### Phage isolation and routine phage culturing

Raw sewage was acquired from the Finham Treatment Works, Coventry, United Kingdom (coordinates 52.36, -1.51). The sediment was removed through centrifugation at 3000 *g*, 4°C, 10 minutes, and the supernatant filtered through a 0.22 µm filter (Cytiva, USA). The supernatant was combined with mid-exponential phase *B. cenocepacia* K56-2 wildtype in 2X LB media and incubated for 18 hours at 37°C with agitation. Following the coincubation, bacterial cells were pelleted (3000 *g*, 10 minutes, room temperature), and the supernatant was filtered (0.22 µm), diluted in SM buffer (100 mM NaCl, 8 mM MgSO_4_, 10 mM Tris pH 7.5), and plated in a soft agar overlay. The plates were incubated at 37°C for 24 hours. Plaques were picked using a P1000 tip, resuspended in 100 µL SM buffer, vortexed, and the samples diluted and soft agar overlay plates were made. Isolation was repeated for four generations until a consistent plaque morphology was achieved.

Phage were routinely cultured in LB supplemented with 2 mM CaCl_2_ + 2 mM MgSO_4_ unless otherwise specified. For soft agar overlay plates 100 µL of overnight bacterial culture and 100 µL of phage were combined with 3 mL of 0.5% soft agar. Phage stocks were made as previously described^53^. Briefly, fully lysed plaque plates were flooded with 5 mL SM buffer and left to adsorb for 30 minutes at room temperature. The soft agar layer was scraped off using a disposable spreader, collected and the samples were agitated to further break down the agar and release the phage. The agar was pelleted at 3000 *g* for 30 minutes at room temperature, supernatant filtered through a 0.22 µm filter (Cytiva, USA) and the stock was stored at 4°C and with 25% glycerol at -70°C.

### DNA isolation, long- and short-read sequencing, and genome assembly

Phage DNA was purified from stocks >1*10^9^ pfu/mL using a phenol:chloroform extraction as outlined in the PCI/SDS DNA Extraction protocol at PhagesDB^54^. The purified DNA was submitted to MicrobesNG (Birmingham, UK) for paired-end Illumina sequencing and Oxford Nanopore Technologies (ONT) long-read sequencing, and to NU-OMICS (Newcastle, UK) for PacBio sequencing. The PacBio library was prepared using PacBio HiFi Plex Prep Kit (PacBio, USA). Briefly, samples were quantified using the Qubit 1X DNA High Sensitivity Assay Kit (Invitrogen, USA) and then normalised to a concentration of 300 ng in 50 µl. DNA was sheared to 7-10 kb size range using the QIAGEN TissueLyzer II 2x96 well 2 ml plate adapter with a cycle of 15 Hz for 5 minutes. Sheared DNA was cleaned using 1x SMRTbell cleanup beads and prepared according to manufacturer’s instructions. The final library was quantified using the Qubit High Sensitivity dsDNA Assay Kit (Invitrogen, USA), diluted to 300 pM, spiked with control DNA and loaded on to the PacBio Revio with the movie length set to 24 hours. Data was demultiplexed and exported in fastq format using SMRTLink (v.25.3).

Reads were trimmed with Cutadapt v4.6^55^ and a hybrid assembly was achieved using SPAdes genome assembler v3.15.5^56^. The assembly map was visualised using Bandage^57^ and the three connected contigs assembled into a single sequence using the overlapping sequence. Reads were mapped to the reference genome using minimap2^58^, the alignments processed using SAMtools^59^ and the nucleic acid read mapping visualised using Qualimap2^60^ and Integrative Genome Viewer (IGV)^61^. The final genome sequence was manually curated to match the physical ends of the repeat sequences. The sequence quality and identification of linear form was undertaken using QUAST v5.2.0^62^.

### Genome annotation and phylogenetic tree generation

The genome was annotated using Pharokka v1.7.1^63^ that encompasses PHANOTATE for coding sequence (CDS) prediction^64^, tRNAscan-SE 2.0 for tRNA prediction^65^, and CRT for CRISPR prediction^66^. Functional annotation was performed using MMseqs2^67^ and PyHMMER^68^ to match each CDS to the VFDB^69^, PHROG^70^, and CARD^71^ databases. Contigs were matched to their closest relatives in the INPHARED database^72^ using mash^73^. Additional functional annotation was undertaken using PHOLD^74^. Genome plots were generated with pyCirclize^75^. Taxonomy was assigned using TaxMyPhage^76^ with the distance increased to 0.3. The phage lifestyle was determined using PhageLeads^77^. Proteome sequences predicted from annotated genes were made using Clinker^78^ to the closest relatives: *Burkholderia* phage BcepSaruman (MK552140) and *Burkholderia* phage BcepSauron (MK552140). The phylogenetic tree was generated using VIPtree v1.1.2^79^ using the INPHARED database^72^ and visualised using iTOL^80^. Phage encoded anti-defence systems were identified with DefenseFinder^81–84^.

### Polymerase Chain Reaction (PCR)

For standard PCR PCRBIO Taq Mix Red (PCR Biosystems Ltd, UK) was used according to manufacturer’s instructions using purified virion DNA. When plaques were used as the source of DNA the plaque was picked using a P10 pipette tip and the phage resuspended in 20 µL of sterile water. The initial lysis cycle was extended to 10 minutes to allow for virion lysis. For PCR of large fragments (>10 kb) Q5® High-Fidelity DNA polymerase mastermix (New England Biolabs, USA) was used as per manufacturer’s instructions. The DNA was separated in a 1% (w/v) agarose gel containing SYBR Safe DNA Stain (Thermo Fisher Scientific, USA) and visualised using the GeneGenius Bio Imaging System (Syngene, UK). All primers were synthesised by Integrated DNA Technologies (Iowa, USA) and are listed in Supplementary Table 2. DNA fragments were purified using the Monarch PCR & DNA or Gel extraction kits (New England Biolabs, USA). Amplicon sequences were checked by Sanger sequencing (Eurofins Genomics, UK).

### Pulsed-Field Gel Electrophoresis (PFGE)

PFGE was performed as previously described^85^. A high titre phage lysate (9x10^9^ pfu/mL) was mixed with equal volumes of SeaKem Gold Agarose (Lonza Bioscience, Switzerland) to create agarose plugs. Plugs were incubated in lysis buffer (100 mM EDTA, 100 mM Tris-HCl pH 8.0, 1% SDS, 0.5 mg/mL Proteinase K) at 55°C overnight. Plugs were washed thrice in 0.5X TBE buffer (45 mM Tris borate and 1 mM EDTA) and loaded into a 1% (w/v) SeaKem Gold Agarose gel with Lambda and yeast chromosome PFGE markers (Sigma-Aldrich, USA). Electrophoresis was conducted using a CHEF Mapper® XA Pulsed Field Electrophoresis System (Bio-Rad, USA) Auto Algorithm at 6 V/cm for 22 hours at 14°C. The gel was post-stained with SYBR Safe DNA stain and visualised using the GeneGenius Bio Imaging System (Syngene, UK).

### Plaque morphology

Images of plaques following incubation at 30°C for 24 hours were acquired using a CanoScan LiDE 400 scanner (Canon, Japan). Plaque sizes were determined using ImageJ^86^.

### Transmission electron microscopy (TEM)

For TEM, high-titre phage lysate was buffer exchanged into SM buffer using Vivaspin 20 centrifugal concentrator 300 kDa MWCO columns (Cytiva, USA). A 5 µL drop of phage was briefly placed on a glow-discharged carbon/formvar coated 300 mesh copper TEM grid (EM Resolutions, UK) and the grids were stained with 2% (w/v) uranyl acetate for 1 minute. Images were acquired on the JEOL 2100 Plus electron microscope (JEOL, Japan) with a Gatan ultrascan camera (Gatan, USA). The virion size was determined using ImageJ^86^.

### Phage killing kinetics

Wildtype *B. cenocepacia* K56-2 in biological triplicate were grown to mid-exponential and diluted to OD_600nm_ 0.001 (4x10^6^ colony forming units (cfu)/mL) in LB + 2mM MgSO_4_ + 2mM CaCl_2_ and combined with φMatilda at Multiplicity of Infection (MOI) 225, MOI 100, and subsequent ten-fold dilutions to a final volume of 200 µL in a 96-well plate covered with a Breathe-Easy® membrane (Sigma-Aldrich, USA). Plates were incubated in a Spark Multimode Microplate reader (Tecan, Switzerland) for 24 hours at 30°C or 37°C with constant agitation (180 rpm). Measurements were taken every 20 minutes at OD_600nm_. The virulence index (V_i_) and growth reduction coefficient (GRC) were determined using the equation 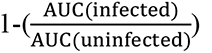 where the area under the curve (AUC) was calculated using gcplyr^87^ in R^88^. For the V_i_ only the AUC for the lag and exponential phases was used, while the GRC utilised the AUC of the entire curve.

### Host range analysis and efficiency of plating

Dilutions of phage lysates (starting 5x10^9^ pfu/mL) were spotted (10 µL) onto soft agar overlay plates containing 100 µL of stationary phase bacteria (Supplementary Table 1) and activity was assessed after 48 hours at 30°C. The efficiency of plating (EOP) from the biological triplicate of the susceptible strains was determined by spotting 10 µL of phage lysate diluted in SM buffer in technical triplicate onto a soft agar overlay plate and incubated at 30°C for 36 hours. Plaques were quantified and the EOP relative to the *B. cenocepacia* K56-2 strain was determined using the equation 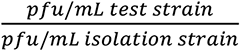. For complemented *B. cenocepacia* K56-2 mutants, 100 µg/mL trimethoprim and 0.1% L-rhamnose were supplemented into the soft agar and the EOP was compared to the *B. cenocepacia* K56-2 + pSCrhaB2 strain as above. For quantification purposes, when clearance was observed at dilutions above the 5*10^9^ pfu/mL stock but individual plaques could not be counted, a plaque count of 1 pfu was assumed at the highest dilution showing clearance.

### Adsorption assay

Mid-exponential wildtype *B. cenocepacia* K56-2 (OD_600nm_ 0.3, 3x10^8^ cfu/mL) was combined with φMatilda at a MOI of 0.1 and cultures incubated at 30°C static. Samples were taken every 5 minutes for 20 minutes, centrifuged at 13,000 rpm at room temperature, for 2 minutes to remove the bacteria, and the supernatant was immediately diluted in SM buffer. Phages were quantified through 10 µL spots in technical duplicate plated on soft agar overlay plates containing wildtype *B. cenocepacia* K56-2. The assay was repeated in biological quadruplicate. To assess the adsorption of φMatilda to the flagella mutant strains, mid-exponential cultures (*B. cenocepacia* wildtype, Δ*fliF*, and Δ*motB*) were diluted to OD_600nm_ 0.3 and combined with Matilda at an MOI of 0.01. Cultures were incubated for 10 minutes, static, and samples pelleted to remove any bound phages and the unbound phages were serially diluted in SM buffer and quantified on soft agar overlay plates containing wildtype *B. cenocepacia* K56-2. The assay was repeated in biological triplicate.

### One-step growth curve

Early exponential phase (OD_600nm_ 0.1) wildtype *B. cenocepacia* K56-2 cells in biological quadruplicate were infected with phage at an MOI of 0.1 or an MOI of 3 in 10 mL. The phages were adsorbed to the bacteria for 10 minutes at 30°C, static. Bacterial cells were pelleted at 10,000 *g*, room temperature, 10 minutes, and washed once with SM buffer, 13,000 *g*, room temperature, 2 minutes, to remove unbound phage. The pellets were resuspended in 10 mL LB +2 mM CaCl_2_ + 2 mM MgSO_4_. Samples were incubated at 30°C, 150 rpm for 120 minutes with 200 µL of cells harvested every 10 minutes following the pellet resuspension (Time 0). Harvested cells were immediately spun at 13,000 *g* for 2 minutes, room temperature, and the supernatant serially diluted and plated in soft agar overlay plates.

### Liquid chromatography nucleoside analysis method on phage DNA

The nucleoside composition of phage DNA was analysed using high-performance liquid chromatography-mass spectrometry (HPLC-MS) on nuclease- and phosphatase-digested DNA. Approximately 1-2 µg of phage DNA (φMatilda virion DNA or reference DNA from phage λ (New England Biolabs, Cat# N3011)) was treated with the Nucleoside Digestion Mix (New England Biolabs, Cat# M0649) following the manufacturer’s protocol at 37°C for >2h. The resulting nucleosides mixture was filtered through a hydrophilic PTFE 0.2 µm centrifugal filter, and the filtrate was subjected to reverse-phase HPLC for nucleoside separation and detection. LC/MS was performed on an Agilent LC-MSD XT system configured with a 1290 Infinity II LC system, a G7117 Diode Array Detector, and a G6135 single quadrupole mass detector. A Waters Atlantis T3 C18 column (100Å, 3 µm, 4.6 × 150 mm) was used for chromatography with the instrument, which was operated at a flow rate of 0.6 mL/min with a binary linear gradient of mobile phases consisting of 10 mM ammonium acetate (pH 4.5) and methanol over 6 min. We monitored the chromatographic course at 260 nm. Agilent ChemStation software was used for primary LC–MS data processing. The yielded chromatograms were further rendered and annotated in Adobe Illustrator. ChemStation’s integration function quantified the peak area of each nucleoside species resolved in the HPLC trace. The peak areas were divided by the corresponding nucleoside molar extinction coefficient (ε) at 260 nm. The extinction coefficients used were 7100 cm^-1^ M^-1^ for dC, 15066 cm^-1^ M^-1^ for dG, 8560 cm^-1^ M^-1^ for dT, and 15060 cm-1 M^-1^ for dA^89^. The percent replacement of canonical bases with the hypermodified base for each phage is calculated from the (ε)-adjusted quantity of nucleoside.

### Sample preparation for transcriptomics and proteomics

Early exponential *B. cenocepacia* K56-2 cells (OD_600nm_ 0.1) were prepared in biological quadruplicate and samples split into two sets. One set of replicates was infected with φMatilda at an MOI of 3 and the other set was treated with an equal volume of SM buffer (uninfected). Samples were placed at 30°C for 10 minutes to allow the phage to adsorb, and then pelleted at 3,000 *g*, 4°C for 10 minutes, the supernatant discarded, and the cells resuspended in LB +2 mM CaCl_2_ + 2 mM MgSO_4_. An aliquot of cells was harvested immediately following resuspension, and all samples were incubated at 30°C for 20 minutes with additional cell harvests at 10 and 20 minutes. Harvested cells were immediately pelleted at 13,000 *g*, 2 minutes, room temperature, and washed once with PBS. The supernatant was discarded and the cell pellets immediately flash frozen in an ethanol dry ice bath and stored at -70°C. Samples for proteomic and transcriptomic analysis were prepared independently.

### Transcriptomics

Frozen cell pellets were sent to Genewiz (Azenta Life Sciences, UK) for RNA extraction and rRNA depletion. Libraries were prepared for paired-end sequencing and run on a NovaSeq X Plus (Illumina, USA) to a target depth of approximately 20 million reads per sample.

### Transcriptomic data analysis

Reads were trimmed using Cutadapt v2.6^55^ and the sample quality was checked using fastQC v0.12.1^90^. Reads were aligned to the *B. cenocepacia* K56-2 chromosomes and cryptic plasmid (CP053300, CP053301, CP053302, CP053303) and the *Burkholderia* phage Matilda genome (Accession ERS30300387) using bwa-mem v0.7.18^91^. Files were sorted and indexed using SAMtools v1.3.1^59^. Counts were generated using HTSeq v2.0.3^92^ using default parameters except non-unique was set to ‘all’, order set to ‘pos’, and minaqual set to 0 to enable mapping to the repeated sequences. All subsequent analysis was conducted using R v4.5.1^88^. Differentially expressed genes were identified using DESeq2 v1.50.2^93^. Genes with less than 5 reads across three replicates in any sample were removed from analysis. Log2 values were adjusted using the shrinkage method of apeglm v1.32.0^94^. The threshold for differentially expressed genes was defined as adjusted *p*-value ≤0.05 and abs(log2FC) ≥ 1. Sample clustering was assessed by PCA analysis using the regularised-log normalised log2 values. The optimal number of expression profile clusters was determined using the silhouette cluster method using factoextra v2.0.0 ^95^and clusters were determined using pheatmap v1.0.13^96^. Data was normalised to transcripts per million (TPM).

### Data-dependent acquisition mass spectrometry

Cell pellets were resuspended in 4% SDS. 100 mM Tris pH 7.5 and lysed by boiling twice at 99°C for 10 minutes, 2000 rpm on an Eppendorf ThermoMixer©. Protein concentrations assessed using bicinchoninic acid protein assays (Thermo Fisher Scientific, USA) and 100 μg of each sample was precipitated in 4X ice-cold acetone at -20°C for 18 hours, and centrifuged at 3,300 *g*, 0°C, 10 minutes. The supernatant was discarded and the proteins were resuspended in water at room temperature. The acetone precipitation was repeated with an incubation of 4 hours at -20°C, and the precipitated proteins were prepared for digestion using S-trap mini columns (Protifi, USA) according to the manufacturer’s instructions. Briefly, samples were reduced with 10 mM dithiothreitol for 10 mins at 95 °C and then alkylated with 40 mM Iodoacetamide in the dark for 1 hour. Samples were acidified to 1.2% phosphoric acid and diluted with seven volumes of S-trap wash buffer (90% methanol, 100 mM Tetraethylammonium bromide pH 7.1) before being loaded onto S-traps and washed 3 times with 400 μL of S-trap wash buffer. Samples were then digested with 4 μg of Trypsin (a 1:50 protease/protein ratio) in 100 mM Tetraethylammonium bromide overnight at 37 °C before being collected by centrifugation with washes of 100 mM Tetraethylammonium bromide, followed by 0.2% formic acid, and then 0.2% formic acid / 50% acetonitrile. Samples were dried down and further cleaned up using C18 Stage^97,98^ tips to ensure the removal of any particulate matter.

C18 cleaned up peptide samples were re-suspended in Buffer A* (2% acetonitrile, 0.1% trifluoroacetic acid in Milli-Q water) and separated using a two-column chromatography set-up on a Dionex Ultimate 3000 UPLC composed of a PepMap100 C18 20 mm x 75 μm trap and a PepMap C18 500 mm x 75 μm analytical column (Thermo Fisher Scientific, USA) coupled to an Orbitrap Fusion™ Eclipse Tribrid™ Mass Spectrometer (Thermo Fisher Scientific, USA). 95-minute gradients were run for each sample, with samples loaded onto the trap column with 98% Buffer A (2% acetonitrile, 0.1% formic acid in Milli-Q water, v/v) and 2% Buffer B (80% acetonitrile, 0.1% formic acid, v/v) for 6 minutes at 5μl/min then peptides separated by altering the buffer composition from 2% Buffer B to 23% B over 59 minutes, then from 23% B to 40% B over 8 minutes, then from 40% B to 80% B over 5 minutes, the composition was held at 80% B for 5 minutes, and then dropped to 2% B over 3 minute and held at 2% B for another 9 minutes. A data-dependent approach was utilised. A single Orbitrap MS scan (375-2000 m/z, maximal injection time of 50 ms, an AGC of maximum of 100% and a resolution of 120k) was acquired every 3 seconds followed by Orbitrap MS/MS HCD scans of precursors (NCE 30%, maximal injection time of 80 ms, an AGC set to a maximum of 300% ions and a resolution of 30k).

### Proteomic data analysis

Samples were searched against the *B. cenocepacia* K56-2 proteome (Uniprot: UP000034659) and an in-house generated proteome of φMatilda (Accession ERS30300387) using MaxQuant (v2.4.7.0)^99^, allowing carbamidomethylation of cysteine set as a fixed modification and oxidation of methionine as a variable modification. Searches were performed with Trypsin cleavage specificity, allowing 2 miscleavage events with a maximum false discovery rate (FDR) of 1.0 % set for protein and peptide identifications. Proteins within the repeat region are only mapped to one of the repeats and are not duplicated. The LFQ and “Match Between Run” options were enabled to allow comparison between samples. The resulting data files were processed using Perseus (v1.4.0.6)^100^. Data from the time course infection were filtered for proteins present in at least 3 out of any 4 samples. Missing values were imputed based on the total observed protein intensities with a range of 0.3 σ and a downshift of 2.1 σ. Statistical analysis of the time course samples was undertaken in Perseus using two-tailed unpaired T-tests.

### Visualisation and statistical analysis

Plots and statistical analyses were generated using R v4.5.1^88^ using ggplot2 v4.0.2^101^. For comparison between two samples (host range EOP) a Mann-Whitney U test was performed to determine any statistically significant differences. For comparisons of changes in the phage killing kinetics assay and phage adsorption, a Kruskal-Wallis test was used.

## Results

### Isolation of novel *Burkholderia*-targeting phage

To expand on the limited pool of characterised phages targeting BCC species, a novel phage was isolated from raw sewage following coincubation with the model strain *B. cenocepacia* K56-2^39^. Following the initial isolation at 37°C, where turbid plaques were observed, the phage was routinely cultured at 30°C, resulting in clearer and more pronounced plaques (0.3-1.6 mm, Figure 1A). To understand the breadth of the host range, the phage was tested against a panel of *Burkholderia cepacia* complex species and shown to be active against 17 of the 28 isolates tested (Figure 1B). This corresponded to activity against 10 BCC species, including *Burkholderia multivorans,* the most common *Burkholderia* species currently infecting people with CF^102^. The virion morphology was visualised using TEM, where a large myovirus-like structure was observed (Figure 1C). The capsid width was 110 nm ± 4.6 nm with a contractile tail of 82 nm ± 5.1 nm in length. Very short tail fibers were observed, and the phage was captured interacting with outer membrane vesicles (OMVs; Figure 1C green arrows), including instances with a contracted tail, suggesting injection into the OMV. This phage was designated *Burkholderia* phage vB_BceM_Matilda (φMatilda). A secondary phage considerably smaller in size than φMatilda was occasionally observed in the TEM. *B. cenocepacia* K56-2 encodes a known prophage, *Burkholderia* phage KS10, which shares similar morphology to this secondary phage. Due to the low abundance and morphological similarity to phage KS10, the secondary phage was likely a natural contaminant and was not considered for further analysis.

**Figure 1:**
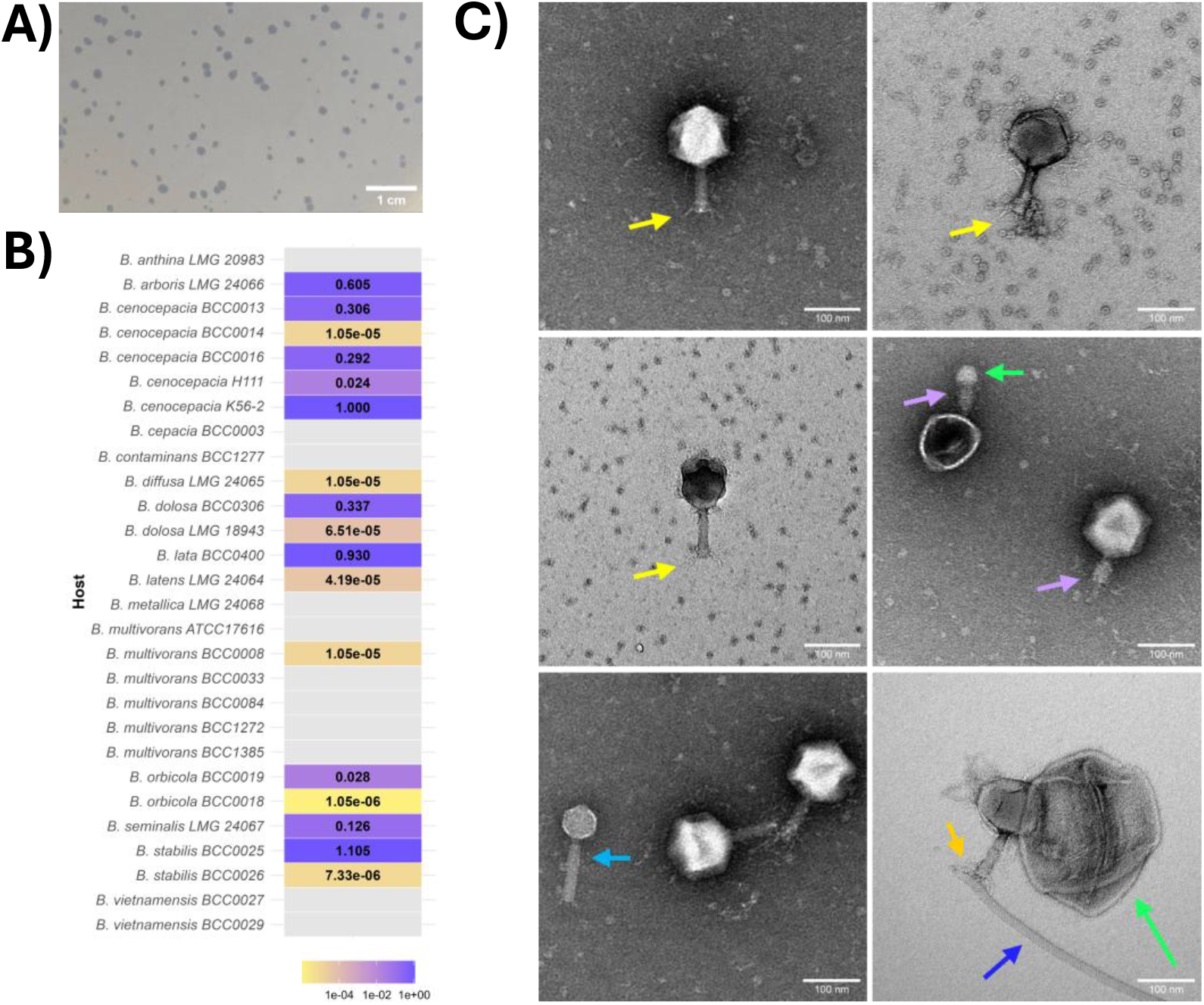
Morphology and host range of φMatilda. **A)** Plaque morphology of φMatilda on a wildtype *B. cenocepacia* K56-2 lawn at 30°C for 22 hours. Scale bar represents 1 cm. **B)** *Burkholderia cepacia* complex (BCC) host range of φMatilda. Non-susceptible strains are shown in grey. The scale of EOP relative to the isolation strain *B. cenocepacia* K56-2 is shown in the scale bar. **C)** Transmission electron micrographs of φMatilda. The phages were stained with 2% uranyl acetate and imaged at 60,000-fold magnification. Coloured arrows indicate different structures: Yellow – tail fibers, purple – contracted tails, green – OMVs, light blue – prophage KS10, dark blue – flagellum, orange – phage tail fibers near flagellum. Scale bars represent 100nm.

### An active flagellum is required for φMatilda to infect bacterial cells

The bacterial flagellum is a common phage receptor, with many flagellotropic phages possessing long appendages that wind around this bacterial element^103^. A subset of flagellotropic phages bind directly to the bacterial flagellum or to glycans decorating the flagellum using short tail fibers^103^. Multiple bacterial flagella were observed in the background of the TEM images, possibly copurified as a result of the phage targeting the flagellum, and in two instances, φMatilda was captured interacting with a flagellum via its tail fibers (one example Figure 1A bottom right). To determine if φMatilda utilised the flagellum for infection, the phage was screened against three *B. cenocepacia* K56-2 deletion mutants: Δ*fliF* (aflagellated), Δ*motB* (paralysed flagella), and Δ*pglL* (glycosylation null). φMatilda was unable to infect either of the *B. cenocepacia* Δ*fliF* and Δ*motB* strains, yet infection was restored upon complementation (Figure 2A), confirming that φMatilda requires the flagellum for infection. The infection of the Δ*pglL* strain was indistinguishable from the wildtype, confirming that when the phage targets the flagellum this interaction is not reliant upon the glycans decorating it. Collectively, this confirms that φMatilda requires an active flagellum to facilitate infection and is therefore a flagellotropic phage.

**Figure 2.**
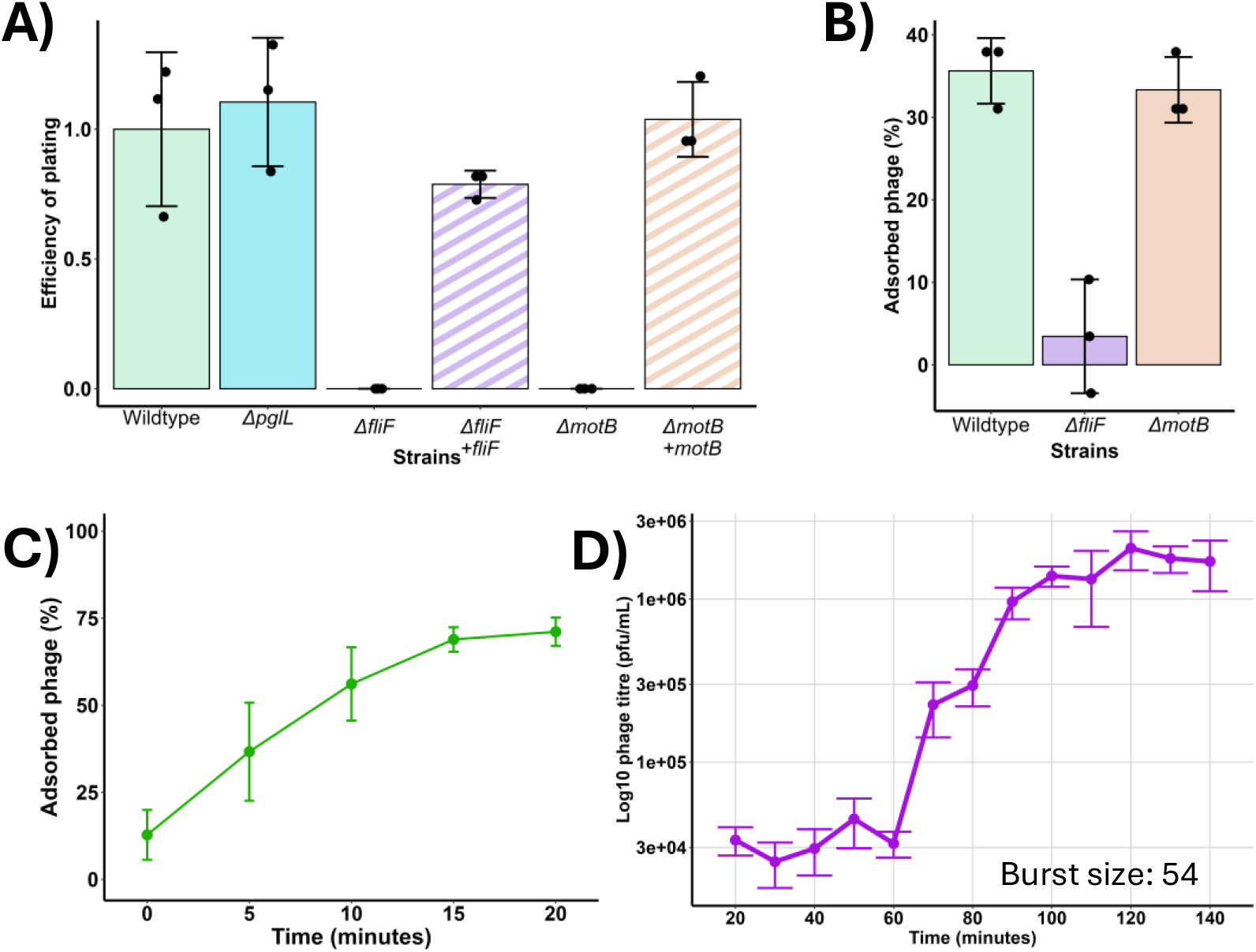
φMatilda receptor and infection dynamics. **A)** EOP of φMatilda against *B. cenocepacia* wildtype and mutant strains: Δ*pglL* – glycosylation null strain, Δ*fliF* – aflagellated strain, Δ*fliF+fliF* – complemented *fliF* mutant, Δ*motB* – paralysed flagella strain, Δ*motB+motB*– complemented *motB* mutant. Data is mean ± SD of three bacterial replicates **B)** Percentage of φMatilda adsorbed to *B. cenocepacia* wildtype, Δ*fliF* and Δ*mot*B strains following an incubation at 30°C for 10 minutes at MOI 0.01. Data is mean ± SD of three bacterial replicates. **C)** Adsorption of φMatilda to wildtype *B. cenocepacia* K56-2 at MOI 0.1 over 20 minutes at 30°C. Percent of adsorbed phage shown. Data is mean ± SD of four bacterial replicates. **D)** One-step growth curve of φMatilda using wildtype *B. cenocepacia* K56-2 at an MOI of 0.1. The mean ± SD of the phage titre at each time point over 120 minutes is shown. N=4. The burst size is indicated on the graph.

As the paralysed and aflagellated strains were resistant to infection, adsorption assays were performed to determine whether the abrogated infection resulted from impaired phage attachment (Figure 2B). Similar levels of adsorption were observed for the wildtype and paralysed flagella (Δ*motB*) strains, consistent with the flagellum being a receptor and the adsorption of φMatilda to the aflagellated strain (*ΔfliF*) was reduced. Together, these results show that φMatilda can bind the paralysed flagellum but cannot proceed to productive infection, whereas the absence of the flagellum largely prohibits phage binding. When φMatilda was tested against only the wildtype strain at an MOI 0.1 over the course of 20 minutes, the proportion of adsorbed phage increased over the first 15 minutes before plateauing at around 75% adsorbed (Figure 2C). To understand the infection dynamics of φMatilda, a one-step growth curve following an adsorption period of 20 minutes at an MOI 0.1 was undertaken. The phage exhibited a latent period of 40 minutes and produced a burst size of 54 virions per bacterial cell (Figure 2D). To understand the differences in how φMatilda controls bacterial growth between 30°C and 37°C, a phage killing kinetic assay was performed (Supplementary Figure 1A and 1B), representing routine culture and human body temperatures respectively. At both temperatures, φMatilda suppressed growth at all MOIs tested. At 37°C the phage was more effective during the exponential phase (Local Virulence Index, Supplementary Figure 1C), whereas at 30°C φMatilda had a sustained impact across the whole growth cycle (Growth Reduction Coefficient, Supplementary Figure 1D). At both temperatures and at all MOIs tested, phage resistant populations emerged as *B. cenocepacia* K56-2 cells entered stationary phase, however bacterial growth never reached that of the uninfected control (Supplementary Figure 1A/B).

### Hybrid assembly reveals three connected contigs

Purified DNA from φMatilda was submitted for paired-end Illumina short-read sequencing, and PacBio HiFi long-read sequencing. A draft hybrid assembly generated from the PacBio and Illumina sequence assembled into three connected contigs of 1,555 bp, 220,813 bp, and 25,181 bp (Figure 3A). No secondary sequences of high coverage were recovered, indicating a pure sample. However, a 37 kb contig was identified with low sequence coverage (5x) matching the transposable prophage, KS10^104^, encoded by the *B. cenocepacia* K56-2 isolation strain, suggesting that the secondary phage identified in the TEM was KS10 (Figure 1C). Subsequent visual exploration using Bandage^57^ revealed a highly complex, non-linear dumbbell-like topology in the assembly graph, characteristic of a linear genome terminated by distinct sets of inverted repeats. The largest contig (220 kb) encoded many essential components expected of a phage genome and we have termed this region the core segment. The small contig (Segment 1) was 1.5 kb in length and included a short 127 bp inverted repeat sequence at each end, which were shared with the 220 kb segment, leaving 1.3 kb of unique sequence in this segment. These 127 bp elements suggest that this terminal segment is connected bidirectionally to the core segment. Similarly, the large terminal segment (25 kb; Segment 3) shared distinct 127 bp elements with the other end of the core segment, suggesting it too could be bidirectional. PCR using primers that bound approximately 250 bp either side of the expected junction between the segments confirmed that the terminal regions were connected to the core segment in a bidirectional manner (Figure 3B, Supplementary Figure 2). All three segments combined to form a single genome with a total preliminary length of 247,041 bp (Supplementary Figure 3A). As the genome length surpasses 200 kb, φMatilda is classified as a jumbo phage.

**Figure 3:**
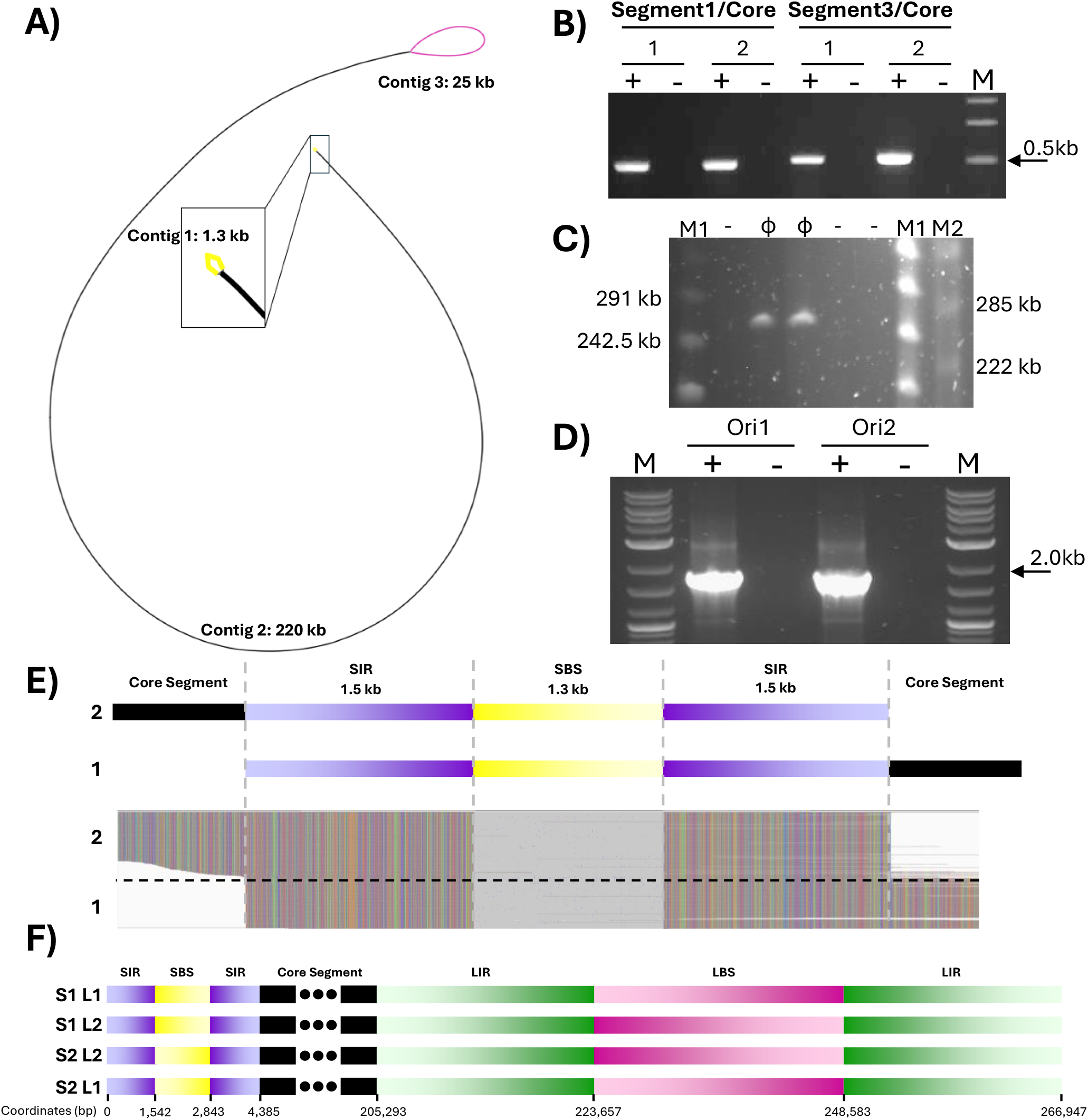
Genome architecture of φMatilda. **A)** Bandage map of the *de novo* assembly of the Matilda genome from the hybrid assembly of Illumina and PacBio sequences. Yellow denotes the short contig (1.3 kb), green denotes the large contig (25 kb), black denotes the core contig (220 kb) **B)** PCR products demonstrating that the bidirectional segments are physically connected to the core segment. Primers AS614/AS616 and AS615/AS616 amplify approximately 500 bp of the region between Segment 1 and the core for orientations 1 and 2 respectively. Primers AS617/AS618 and AS617/AS619 amplify approximately 500 bp of the region between the core segment and Segment 3 for orientations 1 and 2 respectively. Primer sets include no DNA (-) sample. M denotes the NEB 1 kb plus ladder with the 0.5 kb band indicated. **C)** PFGE of the Matilda genome with no DNA controls (-). M1 is the Lambda PFGE ladder with the 291 and 242.5 kb bands marked. M2 is the *S. cerevisiae* PFGE ladder with the 285 kb and 222 kb bands marked. **D)** PCR products amplified from within the SBS to the core segment downstream of the inverted repeats in both orientations. Orientation 1 (Ori1) amplified with AS614/AS643 and Orientation 2 (Ori2) amplified with AS615/AS643. Primer sets amplify a product of approximately 1.9 kb. A no DNA control is present for each primer set (-). M denotes the NEB 1 kb Plus marker with the 2 kb band identified. **E)** Schematic of the SBS end of the φMatilda genome in orientations 1 or 2 (shown on the left). The SBS (yellow) is flanked by the inverted repeats (purple) and connected to the core segment (black) on the left (Orientation 2) or the right (Orientation 1). PacBio reads mapped to the SBS (grey) in a fixed orientation are shown below. Soft clipped sequences are represented by the coloured sequences corresponding to nucleotide base (G – orange, C – blue, A – green, T – red). Orientation 1 finishes with a physical end to the sequence left of the SBS/SIR. Orientation 2 finishes with a physical end to the right of the SBS/SIR. **F)** Schematic of the full φMatilda genome structure showing the four possible genome isomers. SIR and LIR are shown in purple and pink respectively. The SBS and LBS are shown in yellow and green respectively, with the gradient representing the orientation. The orientation of the bidirectional ends is also denoted with **1** and **2** at the start of each genome, with **S** representing SBS and **L** representing LBS.

### Read mapping identified inverted repeat sequences flanking the terminal segments

To reconstruct the true physical sequence of the jumbo phage genome, manual scaffolding was performed by integrating node depth with long read validation. When Illumina and PacBio reads were mapped against the 247 kb sequence, two large coverage peaks were observed (Supplementary Figures 3B and 3C respectively), indicating increased reads in these regions. These peaks occurred at the termini of the core segment sequence (indicated in the purple and pink) rather than within the two terminal segments (Segments 1 and 3). As coverage peaks can indicate repeat structures^13,105^, these regions were scrutinised further, and partially mapped reads were identified. Where the core genome meets the terminal segments, reads that span these regions map correctly to the core genome, but some mapped in an inverted orientation to the terminal segments (Supplementary Figure 3D and 3E). These peak regions were found to form inverted repeats that cap the terminal bidirectional segments (Supplementary Figure 3F). The inverted repeats encompassing the small terminal segment (final size 1,301 bp) were approximately 1.5 kb in length. In contrast the repeats encompassing the large terminal segment (final size 24,927 bp) were 18.3 kb in length. These two repeat sequences are distinct inverted repeat sequences and share no detectable sequence similarity. The features involving the small terminal segment (Segment 1) have been called small bidirectional segment (SBS) and short inverted repeats (SIR). The features involving the large terminal segment (Segment 3) have been called large bidirectional Segment (LBS) and long inverted repeats (LIR). Collectively, the SBS enclosed by paired SIRs and the LBS enclosed by paired LIRs form two distinct terminal architectural elements that bookend the core genome. The bidirectional nature of these terminal elements suggested that alternative genome configurations could exist within the phage population.

### Incorporation of inverted repeats expands the genome size to approximately 267 kb

Incorporating the terminal repeat sequences increased the genome to a final size of 266,947 bp. Sequencing data identified reads that spanned from the core genome into the SBS and LBS in either orientation, suggesting that these sequences were present in two orientations. To confirm that the full length phage genome met the expanded size, pulsed-field gel electrophoresis was performed on the φMatilda genome directly from the lysed phage capsid. This showed a single band of DNA that ran closer to the 285 kb band than the 242.5 kb band (Figure 3C), supporting that the genome is approximately 267 kb, rather than 247 kb. Given the possible bidirectionality of the terminal architectural elements, the single DNA band observed by PFGE may be composed of a mixed population of structural genome isomers that are indistinguishable in size. To understand if the incorporation of the inverted repeats resolved the unusual peaks in the read maps, the Illumina and PacBio reads were remapped onto the full-length 267kb genome containing the full-length inverted repeats (Supplementary Figure 3G). In contrast to the Illumina sequence where the peaks were resolved, the peaks in the PacBio read map were reduced in intensity and width but were still present at the end of the core genome (Supplementary Figure 3H), likely due to the SBS and LBS being present in two orientations.

### Bidirectionality of the SBS experimentally confirmed

The size of the repeats flanking the LBS prohibits PCR-based confirmation of the connection between the core genome and the LBS. Consequently, the SBS alone was used to determine whether the terminal segments were present in both orientations. When primers binding at the end of the SBS (in either orientation 1 or 2) were used with a primer that bound within the core genome, two strong bands were observed for both orientations (Figure 3D; Supplementary Figure 2B). This confirms what was observed in the PacBio sequencing data, where long reads mapped to the SBS in two orientations, spanning from the core genome through one repeat and SBS and into the second repeat (Figure 3E). No reads were observed spanning both SIR and commencing and terminating within the core genome, confirming that the φMatilda genome always terminates at the SIR. This was supported by QUAST analysis of the PacBio genome data that identified no evidence that the genome can circularise.

### Size constraints limit experimental validation of the long bidirectional segment

For the LBS, no reads were identified that commenced in the core genome and extended into the second LIR (>61 kb). However, numerous PacBio reads were identified that commenced in one LIR, crossed the LBS, and terminated in the second LIR (data not shown), supporting the inferred structure of this region. Attempts to generate longer reads, exceeding 61kb, via ONT sequencing were unsuccessful, potentially due to suspected hypermodified bases. However, HPLC analysis of the nucleoside composition of purified φMatilda DNA failed to detect any modifications (Supplementary Figure 4), although RNA bases were present in the sample relative to the λ phage DNA sample.

### Both orientations of the small bidirectional segment are present within single plaques

To determine whether the SBS orientation differed between plaques of varying sizes, the orientation of this segment in fifteen plaques of varying sizes was analysed by PCR (Supplementary Figure 5). All plaques produced bands for both primer sets, confirming that both orientations of the SBS are present within individual plaques regardless of the plaque size. As plaques arise from a single phage virion which would initially possess a single SBS orientation, the presence of both orientations within individual plaques suggests that inversion of the SBS can occur during replication. This structural variation suggests that the terminal segments can undergo intra-molecular homologous recombination across their respective inverted repeats. These findings are consistent with a recombination-mediated “flip-flop” mechanism, which would give rise to a mixed phage population of alternative structural genome isomers, resulting in four possible genome configurations (Figure 3F). Consequently, the final consensus genome was oriented and linearised using a single representative genome orientation.

### φMatilda is representative of a distinct lineage of jumbo phages

As the inverted repeats and bidirectional segments observed in the φMatilda genome have not been previously documented, the placement of this novel phage within phage taxonomy was investigated using TaxMyPhage^76^. Although an initial search with the default distance of 0.2 did not identify any relatives, increasing the MASH distance threshold to 0.3 revealed two related phages from the Sarumanvirus genus, *Burkholderia* phage BcepSaruman (BcepSaruman, 263,735 bp) and *Burkholderia* phage BcepSauron (BcepSauron, 262,653 bp). These two jumbo phages share an average nucleotide identity (ANI) of 93% with each other, but only 36.4% and 36.3% ANI with φMatilda, respectively. This low level of nucleotide identity is consistent with φMatilda representing a distinct lineage at the genus level. A novel genus, Tillivirus, is therefore proposed.

### Comparative genomics reveals a distinct group of φMatilda-related jumbo phages

φMatilda was compared against representative dsDNA phages in the INPHARED dataset and observed to cluster with a group of other jumbo phages (Figure 4A, green star). This group of jumbo phages is very distant from the nucleus-forming jumbo phages (including φKZ^26,106^) (Figure 4A, denoted by the orange circle). In the cluster with φMatilda are BcepSaruman, BcepSauron, and other jumbo phages targeting Gram negative bacteria (Figure 4B). After the Sarumanvirus phages, the next closest relative is the *Ralstonia* phage φRSL1^107,108^. Unlike φMatilda, φRSL1 has a circular genome lacking repeat elements^108^. This fundamental difference in genome architecture suggests that these phages represent distinct families. Conversely, BcepSaruman and BcepSauron have been reported to contain bidirectional elements at either end of a large genome segment^35^, although the size of these elements is not specified. Comparison of the protein sequences of BcepSaruman, BcepSauron, and φMatilda genomes (Supplementary Figure 6) revealed that elements of the φMatilda LBS were present in the Sarumanvirus phages but in the opposite orientation (Supplementary Figure 6 φMatilda green proteins). Notably, the BcepSaruman and BcepSauron genomes commence with a protein corresponding to one located within the SBS in φMatilda, reflecting the absence of an inverted repeat flanking this region. Consistent with this, none of the proteins in the SIR (Figure 3C, Matilda purple proteins) were present in the Sarumanvirus phages, although a homologue of the first protein on the core genome is present in these phages. At the opposite end of the genome, single copy proteins in the BcepSaruman genome have homologues located in the LIRs of φMatilda. While repeat sequences have not been reported in the Sarumanvirus genomes, the presence of shared bidirectional segments suggests that Sarumanvirus and Tillivirus phages represent two distinct genera within the same family, united by their ANI and shared genome architecture.

**Figure 4.**
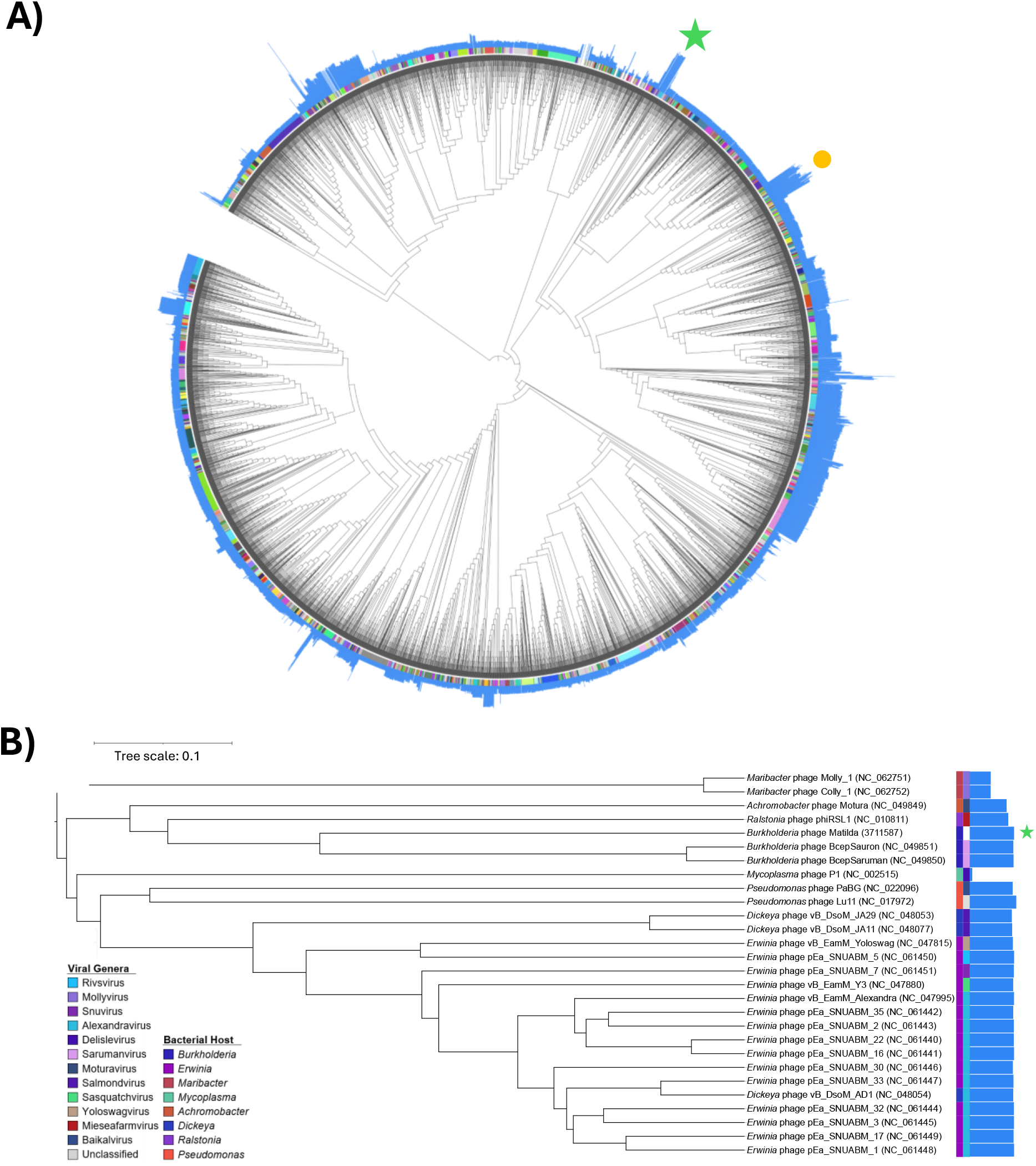
Phylogeny of φMatilda. **A)** Maximum-likelihood (ML) phylogenetic tree of representative dsDNA phages with branch length ignored. The inner coloured ring represents phage genera. The blue outer ring indicates genome length. The jumbo phage cluster containing φMatilda is identified with the green star. The nucleus-forming jumbo phage cluster is identified with the orange circle. **B)** Pruned ML phylogenetic tree showing φMatilda (green star) and its closest relatives. The bacterial host (left) and phage genera (middle) are indicated with the coloured rectangles, and the genome length is indicated by the blue bars (right). The tree scale is shown in the top right.

### A genome dominated by hypothetical genes

The genome of φMatilda encodes 491 genes and 13 tRNA sequences (Supplementary Figure 7) with an average GC content of 55%, about 10% below the average GC content of the *B. cenocepacia* host strain (66%)^109^. Sequence- and structural-based homology searches assigned putative functions to 108 proteins, leaving 78% of the predicted genes encoding hypothetical proteins (Supplementary Table 3).

The SIR and the SBS all encode hypothetical proteins ranging between 37-91 aa for the SIR, and 31-161 aa for the SBS. The genes that bridge the SIR and SBS initiate within the repeat sequence, resulting in a protein with a conserved N-terminal and variable C-terminal. Similarly, genes spanning the LIR-LBS junction also initiate within the repeat sequence. However, none of the junction-spanning genes have a predicted function. The LIR sequences encode 41 proteins, of which 38 encode hypothetical proteins with a median length of 107 aa. The three proteins with expected functions include a RecX-like filament modulator involved in hijacking the host RecA-mediated DNA repair mechanism^110^, an ATP-dependent protease, and an RNA ligase and tail fiber protein attachment catalyst, required for tail fiber attachment to the baseplate^111^. For the LBS 59 of the 65 encode hypothetical proteins ranging from 37-442 aa (median 84 aa). Six further proteins had putative functions including one that was annotated as a tape measure protein. However, at only 194 aa, this protein is substantially smaller than what is expected to span the length of the 82 nm phage tail. The additional presence of a lysozyme domain suggests that this protein is likely a phage tail lysozyme. The other five proteins are involved in DNA and RNA metabolism, including a DNA helicase, homing endonuclease, and tRNA-His guanylyltransferase.

In addition to those found in the LIR and LBS, the core genome of φMatilda encodes 40 genes associated with DNA, RNA, and nucleotide metabolism replication (Supplementary Table 3). Two DNA helicase proteins and one Dda-like helicase are scattered across the core genome, bringing the total number of helicase enzymes to four. There are two DNA polymerases encoded in different regions of the genome of vastly different lengths (1059 aa for TILLLY_0235 and 243 aa for TILLY_0333). The smaller of the two is adjacent to the DNA polymerase processivity factor. Three anti-host defence elements were identified in the genome including an RNA ligase involved in negating the CBASS defence system, a three gene cluster encoding genes associated with the Nicotinamide Adenine Dinucleotide Reconstitution Pathway (NADRP) (TILLY_0129-0131), and a DarB-like antirestriction protein important for blocking bacterial restriction-modification systems. The phage encodes an additional RNA-polymerase sigma factor that may be important for hijacking the bacterial host. Thirty-seven structural proteins for the capsid and tail region were identified including the large terminase that is encoded between a hypothetical protein and a Holliday junction resolvase. Of the four endolysins encoded independently across the genome only two have functional domains; TILLY_0277 contains lambda-like lysozyme domain, and TILLY_0305 contains a glycoside hydrolase domain that can degrade host cell peptidoglycan. The latter of these two proteins is encoded in a locus shared with a holin and a peptidase. Notably no predicted virulence factors, antimicrobial resistance genes, or CRISPR systems were encoded in the φMatilda genome.

### The bidirectional segments and inverted repeats are utilised during a phage infection

A combined proteomic and transcriptomic approach was utilised to determine whether the architectural features were employed by the phage during infection. To ensure that all bacterial cells were infected, and to account for the adsorption rate of Matilda, infection of wildtype *B. cenocepacia* cells was undertaken at an MOI of 3. A one-step growth curve conducted at this same MOI with a 20 minute adsorption/preparation period revealed that bacterial lysis begins after 20 minutes (Supplementary Figure 8), and consequently samples were harvested for transcriptomic and proteomic analysis at 0, 10, and 20 minutes post adsorption (20, 30, and 40 minutes post phage addition). Uninfected bacterial cells were harvested at the same time points as a control. PCA analysis of both the transcriptomic and proteomic samples revealed good separation between the infected and uninfected samples and between timepoints (Supplementary Figure 9A – Transcriptomics, Supplementary Figure 10A – Proteomics). The transcriptomic data revealed that all 491 phage genes were identified across the course of the infection (Supplementary Table 4 and 5). This confirms that the bidirectional segments and inverted repeat sequences were utilised and transcribed during infection, suggesting a functional role of these features. Due to sampling constraints, early gene expression was not captured, with many genes already highly expressed at the first time point (Figure 5A). The clustering of the phage genes was analysed (Supplementary Figure 9B) and while two clusters captured the most variation, three clusters were selected as this clustering better reflects early, middle, and late gene expression (Figure 5B). All genes within the SIR and SBS followed the expression profile of cluster 2, peaking at time 0 (Figure 5C). Similarly, the majority of genes in the LIR and LBS also reflect the cluster 2 expression profile. However, four genes from the LBS and two genes from the LIR (specifically the gene at the junction of the core genome and inverted repeat and the final gene of the repeat) were part of cluster 3. The four genes within the LIR form an operon and encode two hypothetical proteins, a DNA helicase, and a homing endonuclease, which based on the expression profile may be important later in the infection. Many bacterial proteins were upregulated over the course of the expression (Supplementary Figure 9C-E), although few were downregulated. Of those upregulated, K562_13356 which encodes a ribonucleotide-diphosphate reductase subunit beta protein peaked at 10 minutes post-adsorption and may have been hijacked by φMatilda to aid in its genome replication. Of the 19 defence systems encoded by *B. cenocepacia* K56-2, only two increased in response to the phage infection later in the infection (Supplementary Figure 9F): a SoFic system (K562_30093), and a predicted but uncharacterised system (DMS other; K562_20548). Collectively, these data show that genes within both inverted repeats and bidirectional segments are consistently transcribed during infection.

**Figure 5.**
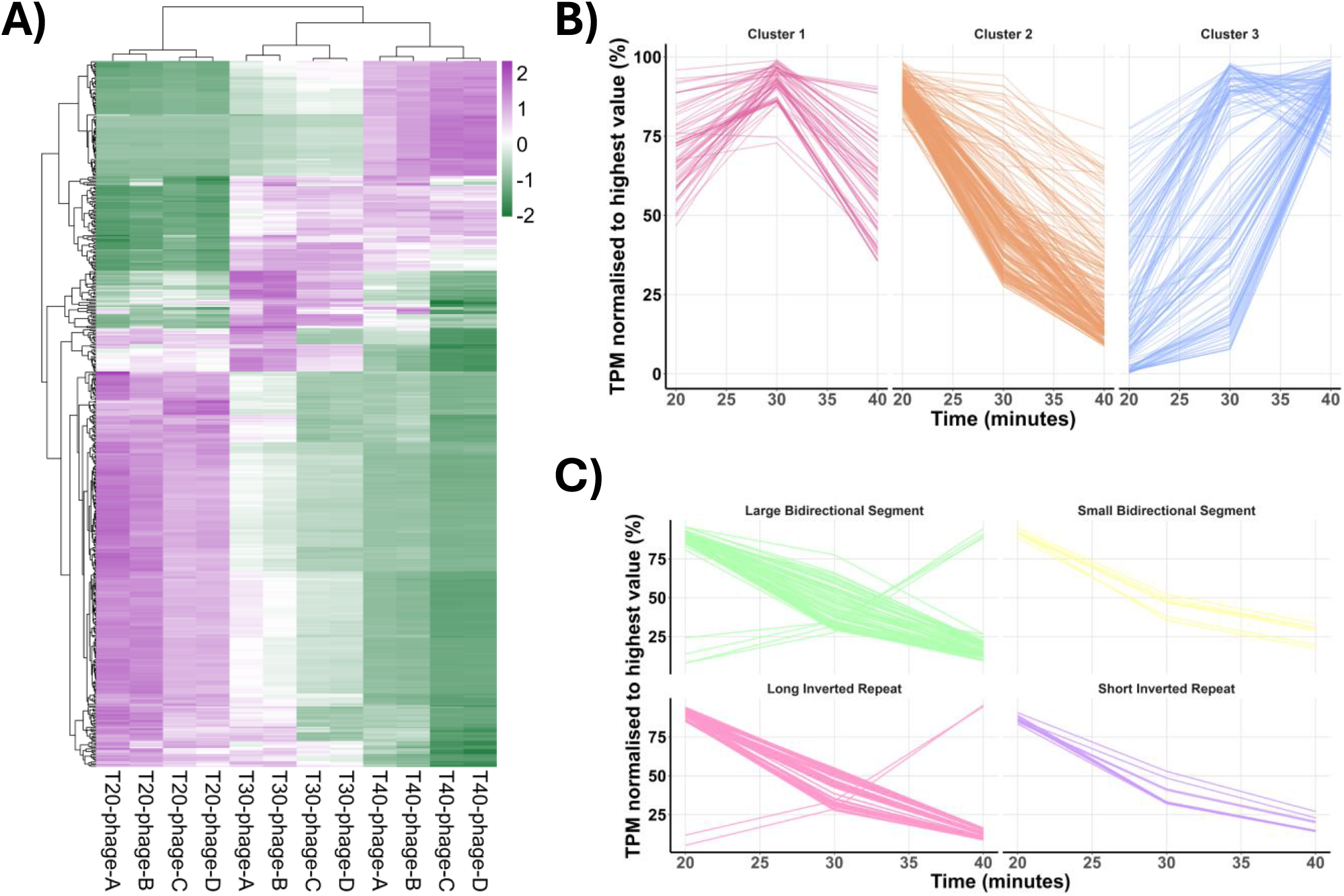
Transcriptomic analysis of φMatilda genes during infection of *B. cenocepacia* K56-2 at 20, 30, and 40 minutes post-adsorption. A) Heatmap of φMatilda gene expression with the individual replicates shown along the bottom of the graph. Hierarchical clustering of phage gene based on expression profiles is shown on the left of the heatmap, with clustering of samples shown at the top. Data represents Z-score normalised TPM values, with the scale shown in the top right. B) Line plot of φMatilda genes separated into three clusters. Cluster 1: Expression peaking at 10 minutes post-adsorption. Cluster 2: Expression peaking at 0 minutes and decreasing over time. Cluster 3: Expression increasing over the course of the infection and peaking at 20 minutes post-adsorption. C) Line plots showing the expression of genes within the SBS (yellow) and LBS (green), and the SIR (purple) and LIR (pink) inverted repeats. Line plot data represent mean TPM values normalised to the highest value observed across the infection.

### Protein response

Proteomic analysis identified 233 φMatilda proteins across the infection time course, capturing the products of just under half of protein-coding genes (Supplementary Table 6). As the infection developed, the protein composition changed (Figure 6A), with lysis and structural protein increasing in abundance towards the later time points. In contrast to the transcriptomics data, fewer proteins were detected at high abundance early in the infection (Figure 6B). A delay between transcription and translation was reflected in the clustering, with many proteins increasing between 0-10 minutes, and a second population increasing between 10-20 minutes, representing the increase in structural and lysis proteins as well as numerous hypotheticals (Figure 6C). The repeat and bidirectional elements showed greater variability in abundance compared to the transcriptional data, where most genes followed a similar expression profile. The only protein identified in the SIR and two proteins encoded in the SBS peaked in abundance in the first time point and were subsequently reduced as the infection progressed (Figure 6D). The majority of the proteins in the LIR and LBS increased in abundance between 0 and 10 minutes and were maintained at a high level. The bacterial proteomic response showed modest early changes, including the increased abundance of two transcriptional regulators (Supplementary Figure 10B). The abundance of these proteins remained elevated at 10 minutes (Supplementary Figure 10C). By 20 minutes post-adsorption, their abundance decreased below levels observed in the uninfected samples, while a glycoside hydrolase became one of the most abundant proteins in the infected samples (Supplementary Figure 10D). Collectively, transcriptomic and proteomic analyses show that genes within the inverted repeats and bidirectional segments are both transcribed and translated during infection, with distinct temporal expression and abundance patterns.

**Figure 6.**
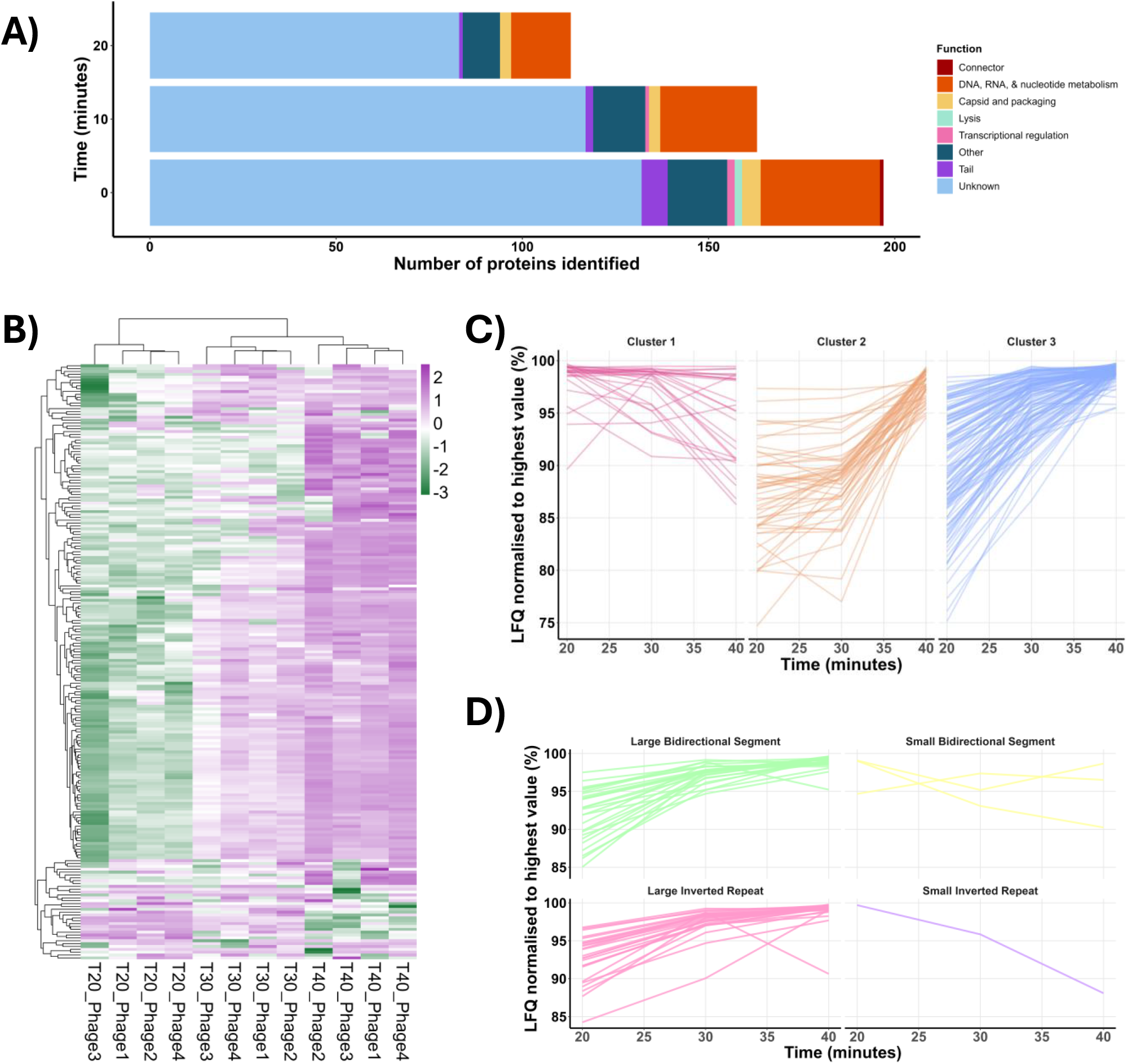
Proteomic analysis of φMatilda genes during infection of *B. cenocepacia* K56-2 at 20, 30, and 40 minutes post-adsorption. **A)** Stacked bar chart showing the total number and functional composition of phage proteins identified at each time point prior to imputation. **B)** Heatmap of φMatilda protein abundance with the individual replicates shown along the bottom of the graph. Hierarchical clustering of phage proteins based on abundance profiles is shown on the left of the heatmap, with clustering of samples shown at the top. Data represents Z-score normalised LFQ values, with the scale shown in the top right. **C**) Line plot of abundance of φMatilda proteins separated into three clusters. Cluster 1: Abundance largely peaking at 0 minutes and decreasing over time. Cluster 2: Abundance increasing over the course of the infection and peaking at 20 minutes post-adsorption. Cluster 3: Abundance peaking at 10 minutes post-adsorption. **D)** Line plots showing the abundance of proteins encoded within the SBS (yellow) and LBS (green), and the SIR (purple) and LIR (pink). Line plot data represent mean LFQ values normalised to the highest value observed across the infection. Proteins from the inverted repeats are only represented once in the data.

## Discussion

This study identified a novel *Burkholderia*-targeting jumbo phage, φMatilda, with an unusual genomic architecture. The linear φMatilda genome has a tripartite structure, in which the terminal regions can associate bidirectionally with the central core. These terminal segments are flanked by long inverted repeat sequences of 1.5 kb and 18 kb that encapsulate bidirectional segments of 1.3 kb and 25 kb respectively. Genes within these distinct architectural elements are expressed during infection.

Repetitive elements are not unusual within phage genomes, with many phages encoding repeat sequences ranging from a few hundred base pairs, such as the 160 bp repeats in T7^112^, to kilobases in length, such as the 10 kb repeats encoded by T5^113^. These repeat sequences are typically direct terminal repeats that bookend the phage genomes and aid in replication^13^. Conversely, the repeats in the φMatilda genome are inverted, and although the genome begins and ends with repeat regions, these terminal sequences differ in both nucleotide composition and length. Inverted repeats have also been documented within other phages with the *Bacillus* phage φ29 encoding short 6 bp inverted repeats at each genome terminus that serve as binding sequences for a DNA capping protein that is required for replication^114,115^. However, these repeats differ from φMatilda by length and context, as the φ29 repeats span the entire genome, rather than defined segments. Additionally, some temperate phages encode invertible DNA sequences that control phage host range, as observed in phages P1 and Mu, which encode 3 kb invertible segments flanked by inverted repeat sequences of ∼600 bp or 34 bp respectively^116–118^. These elements present as a single internal segment, and to date have not been described in lytic phages. Collectively these comparisons highlight the distinct nature of the inverted repeat architecture of φMatilda, relative to characterised phages.

While the common examples of invertible elements do not align with what is observed in φMatilda, similar architecture has been reported in members of the Sarumanvirus genus (BcepSaruman and BcepSauron)^35^. These phages contain a core segment flanked at both termini by distinct bidirectional segments that are physically connected to the core^35^. These phages were isolated from soil in New York, USA more than 20 years prior to the isolation of φMatilda in the United Kingdom, suggesting that this genomic architecture is stable and has been maintained across geographically and temporally diverse isolates. Due to the limited sequencing data available for the Sarumanvirus phages, it is not possible to confirm if they also contain repeat regions. However, such features would be easy to miss, as their identification requires careful manual curation. Similarly, current phage annotation guidelines typically reference phage genomes as arising from a single contig^119,120^. For φMatilda, the terminal segments in the draft assembly encoded mostly hypothetical proteins, whereas the core genome encodes the essential structural proteins (e.g. large terminase, major capsid protein, tail fibers). The presence of essential structural genes within the core genome gives the illusion of a complete genome. Current assembly and annotation workflows are generally optimised to produce a single contiguous genome sequence. Consequently, genomes containing complex terminal architectures such as φMatilda may be inadvertently simplified. This raises the possibility that similar architectures remain underrepresented in current databases and that these additional genomic elements may play an important role in genome organisation, regulation, or stability.

Adeno-associated viruses (AAVs) are small single-stranded DNA eukaryotic viruses that encode short inverted terminal repeats (ITRs)^121^. The ITRs play a crucial role in the stability of the viruses, with shorter ITRs resulting in more stable replication and virus production^122,123^. The genome architecture of AAVs is fundamentally the closest to the genome architecture of φMatilda, with two core differences: φMatilda is a dsDNA genome and therefore could not inherently form the same secondary structure, and 2) The φMatilda ITRs are magnitudes larger than those encoded in AAVs. Despite these differences, it is possible that φMatilda utilises the repeats in a similar way to the ITRs of AAVs to influence genome stability. Additionally, the inverted repeat structures in φMatilda are uniquely associated with bidirectional elements capable of existing in alternative orientations relative to the core genome. Analysis of the small bidirectional segment confirmed the presence of both possible orientations, consistent with the mixed configurations observed in the sequencing data. This observation suggests that the bidirectional segments are not static but may undergo rearrangement during replication or infection. The inverted repeats may facilitate recombination of the segments, or enable the formation of transient circular intermediates, resulting in orientation switching and a dynamic genome architecture.

A core element that remains unknown is what occurs in the bacterial cell immediately following infection. The transcriptomic analysis failed to capture the earliest stages of infection, with many phage genes already highly expressed at the first time point. As a result, it is not possible to determine which phage genes are involved in establishing infection. In the context of genome architecture, these early genes may be involved in genome rearrangement or stabilisation, as observed in φ29^22^, where a terminal capping protein is expressed early to protect the DNA termini. Identifying the protein composition of the φMatilda virion would also have aided in understanding the initial stages of infection, as jumbo phages can carry cargo proteins essential for early infection processes^124–126^. Given that both orientations of the small bidirectional segments were present within individual plaques that likely arose from a single virion, it is possible that genome rearrangement occurs during infection. However, initial population heterogeneity^127^ cannot be excluded. As such, further work is required to elucidate the physical configuration of the φMatilda genome within infected cells. The presence of a linear genome with distinct termini raises questions about how replicated φMatilda genomes are recognised, cleaved, and packaged into the capsid, particularly which terminus serves as the initiation site for genome packaging. Additionally, there is the assumption that the LBS behaves similarly to the SBS. While the sequencing data confirms that both orientations exist, the frequency of the inversion is unknown due to limited experimental validation on this segment. Despite these limitations, the presence of 20 kb of additional repeated DNA encoded within the φMatilda genome represents a substantial increase to the overall genome size. This is expected to increase the metabolic demands of the bacterial cell for both genome replication and production of a larger capsid^23^. The increased requirements for maintaining a larger genome and capsid would not likely be maintained without functional significance. Collectively, these observations highlight the need for further investigation of the mechanistic basis and biological role of the genome architecture of φMatilda, which may represent a novel strategy for genome stability or evasion of bacterial anti-phage defences.

## Conclusion

These results reveal a previously undescribed genome architecture in jumbo phages characterised by bidirectional terminal sequences flanked by inverted repeats. The transcription and translation of these regions during infection supports a functional role of these elements beyond structural purposes. The rearrangement of terminal elements within plaques indicates a dynamic genome configuration, although the benefits of this remain unclear. The conservation of the terminal bidirectional elements in related jumbo phages suggests that this architecture represents a broader, under-recognised genomic strategy, potentially useful for evasion of host defences or genome stability.

## Author contribution

J.M.L, Y.C, T.L, J.J.B, and A.P.S conceived and designed the study. J.M.L isolated and characterised the phage with assistance from H.M and T.M. O.B and J.M.L completed the molecular characterisation of the genome, including PCR and PFGE. Genome assembly was undertaken by J.M.L, O.B, L.B, C.S.M, and R.S, with annotation and analysis by J.M.L. HPLC-MS of phage DNA was performed by YJ.L and P.W. Preparation of samples for transcriptomic and proteomic analysis was performed by H.M and J.M.L. N.E.S undertook the proteomic sample digestion, purification, and data collection, analysis of which was done by J.M.L and H.M. Analysis of transcriptomic samples was completed by J.M.L, L.B, C.S.M, and R.S. A.P.S funded the research. J.M.L drafted the original manuscript and all authors contributed to the review and editing of this manuscript.

## Acknowledgements

We acknowledge funding from the Monash Warwick Alliance AMR training programme to J.M.L. and A.P.S. We also acknowledge support from the BBSRC TRDF (Grant BB/V017217/1) to A.P.S. and from the School of Life Sciences, University of Warwick to O.B.*, H.M.*, T.M.*, J.M.L and A.P.S (*funded by the mBio programme) and support by the National Health & Medical Research Council through an NHMRC Investigator Awards (2016330 to T.L. and 2026130 to J.J.B). E.M. acknowledges funding from the UK CF Trust (CFTP Award No.6) that supported additional curation of *Burkholderia* strains held in the Cardiff UK CF Infection Biorepository (https://cfamr.org.uk/toolkit/uk-cf-infection-biorepository/). We would like to thank Severn Trent Water, Finham Sewage Treatment Works for providing us with the sewage sample. We would like to thank the University of Warwick, School of Life Sciences Media Preparation team for their assistance preparing the media used in this study and Dr Saskia Bakker from the Advanced Bioimaging Research Technology Platform at University of Warwick for her assistance with the transmission electron microscopy. We would like to thank Dr Thomas Naderer from Monash University for his contribution to project design, and Dr Rhys Dunstan at Monash University for his guidance overcoming issues with initial phage isolation. We would like to thank Dr Andrew Millard at the University of Leicester and Prof Jillian Banfield at Monash University/University of California, Berkeley for their assistance deconstructing the distinct read map and mismapped Illumina reads, respectively. We are grateful to Dr Andrew Nelson and Prof Darren Smith at NU-OMICS for undertaking the PacBio sequencing.

## Data Availability

The φMatilda genome sequence is available in the European Nucleotide Archive under study accession PRJEB110049, assembly accession ERS30300387 and taxon ID 3711587. The raw transcriptomic read files are available in the European Nucleotide Archive under study accession PRJEB113199. The proteomics data for the infection experiments has been deposited in the ProteomeXchangeConsortium via the PRIDE partner repository with the data set identifier: PXD070006 Token: T8W1AchBmX00.

---

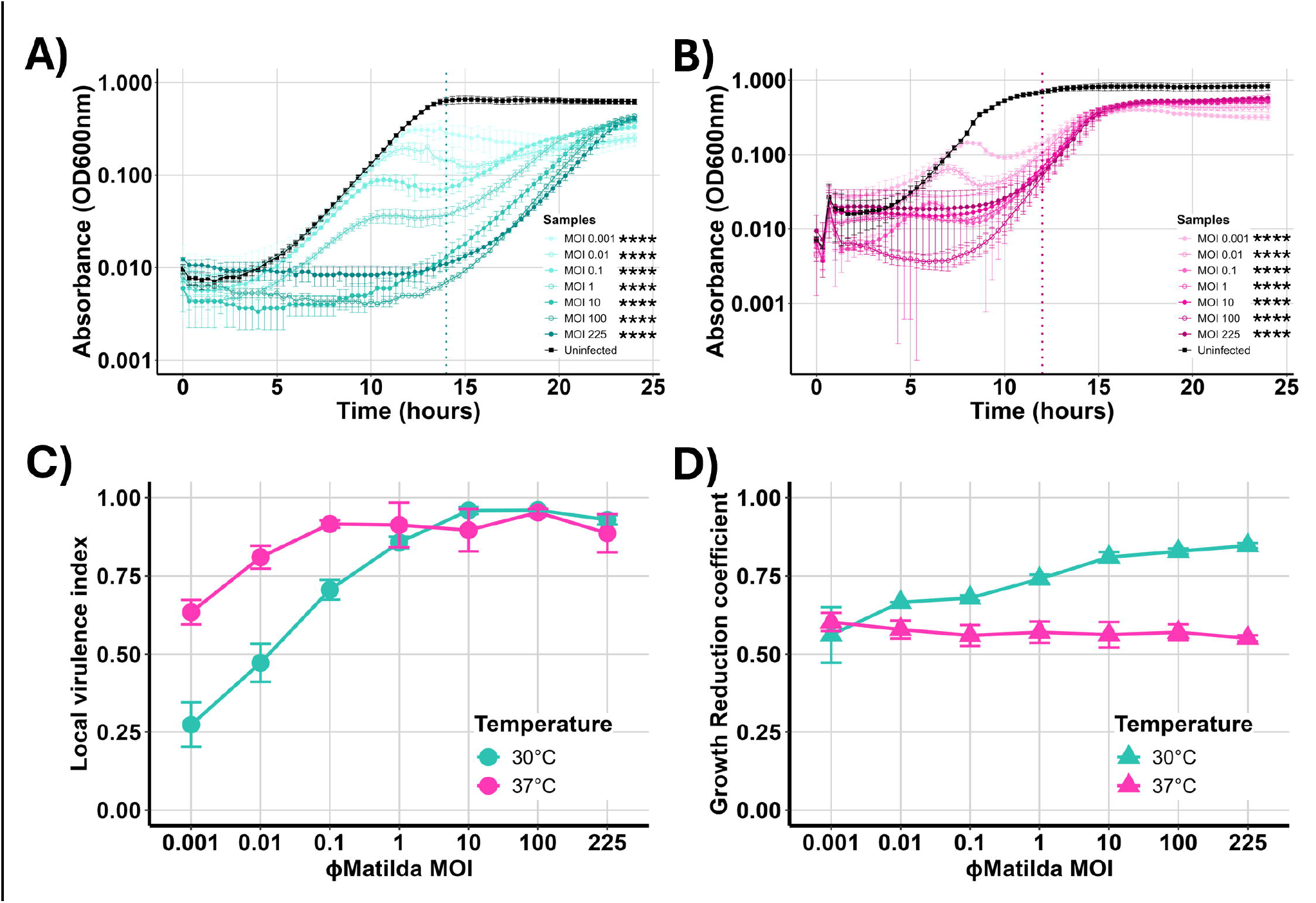

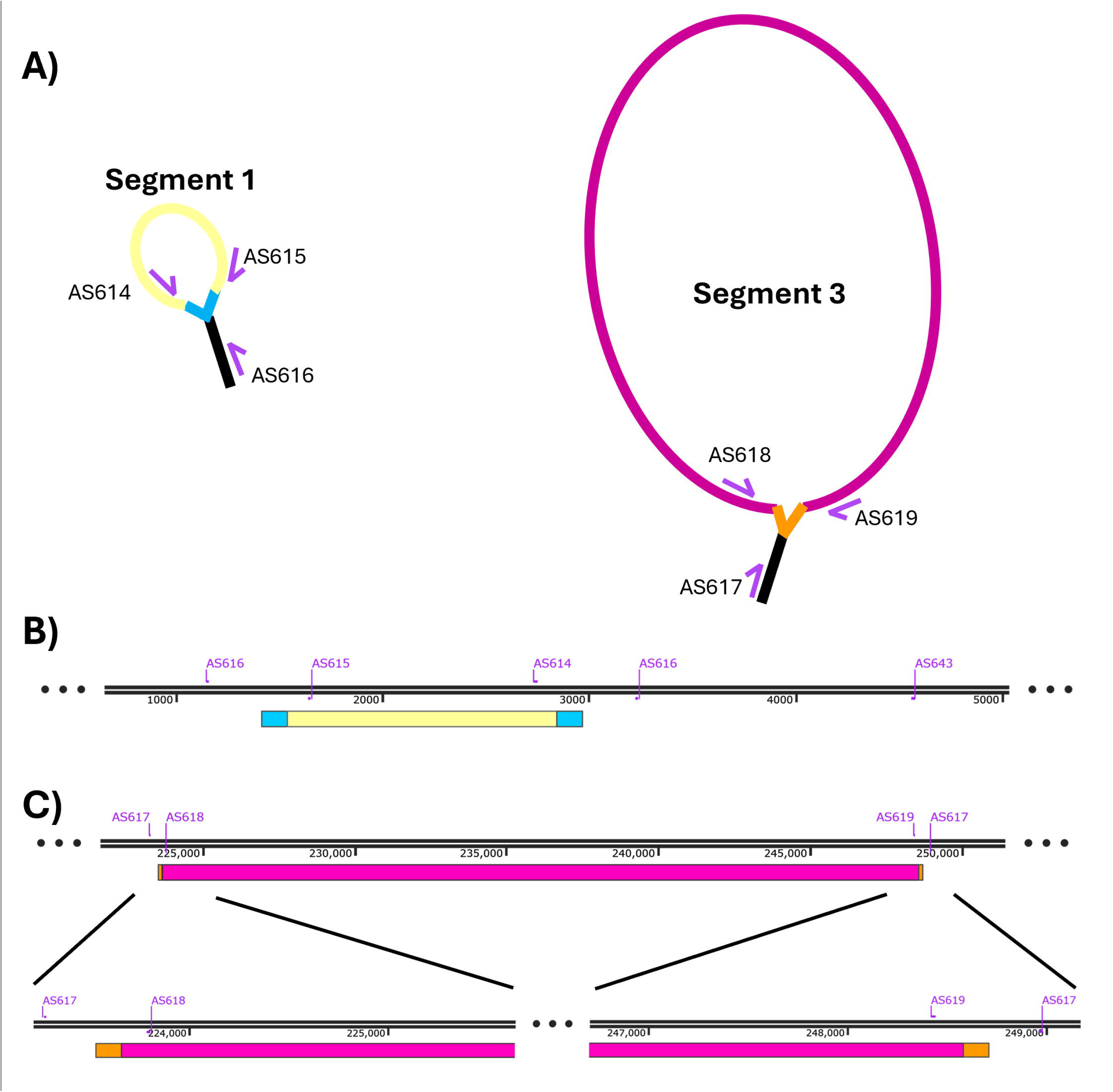

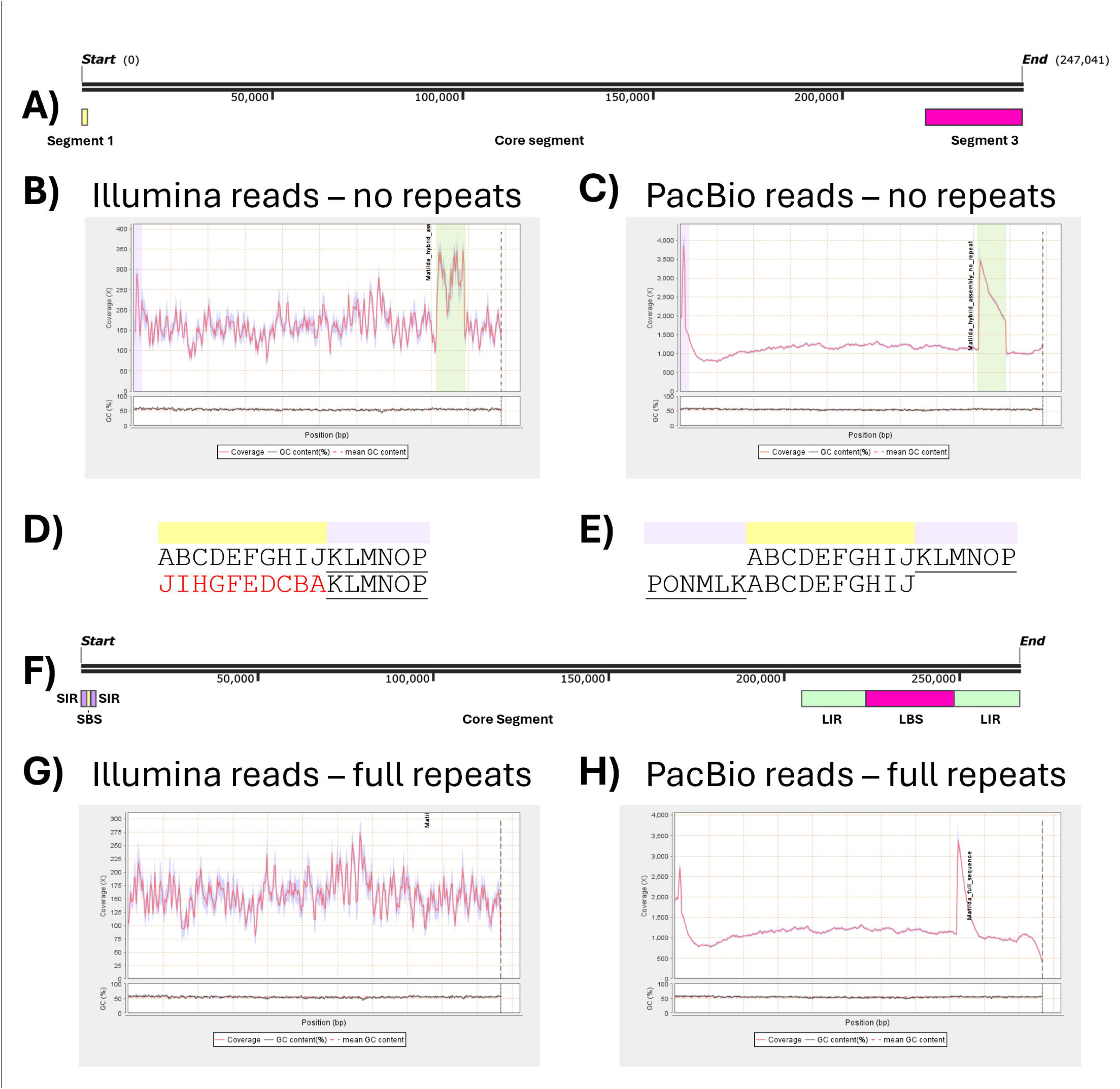

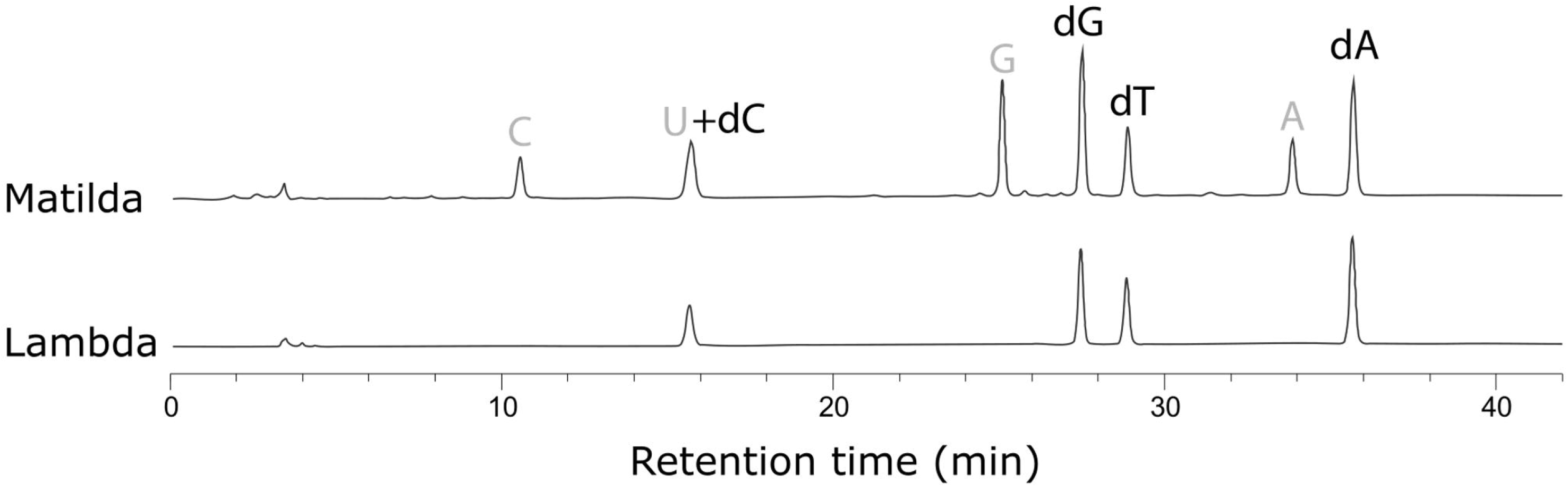

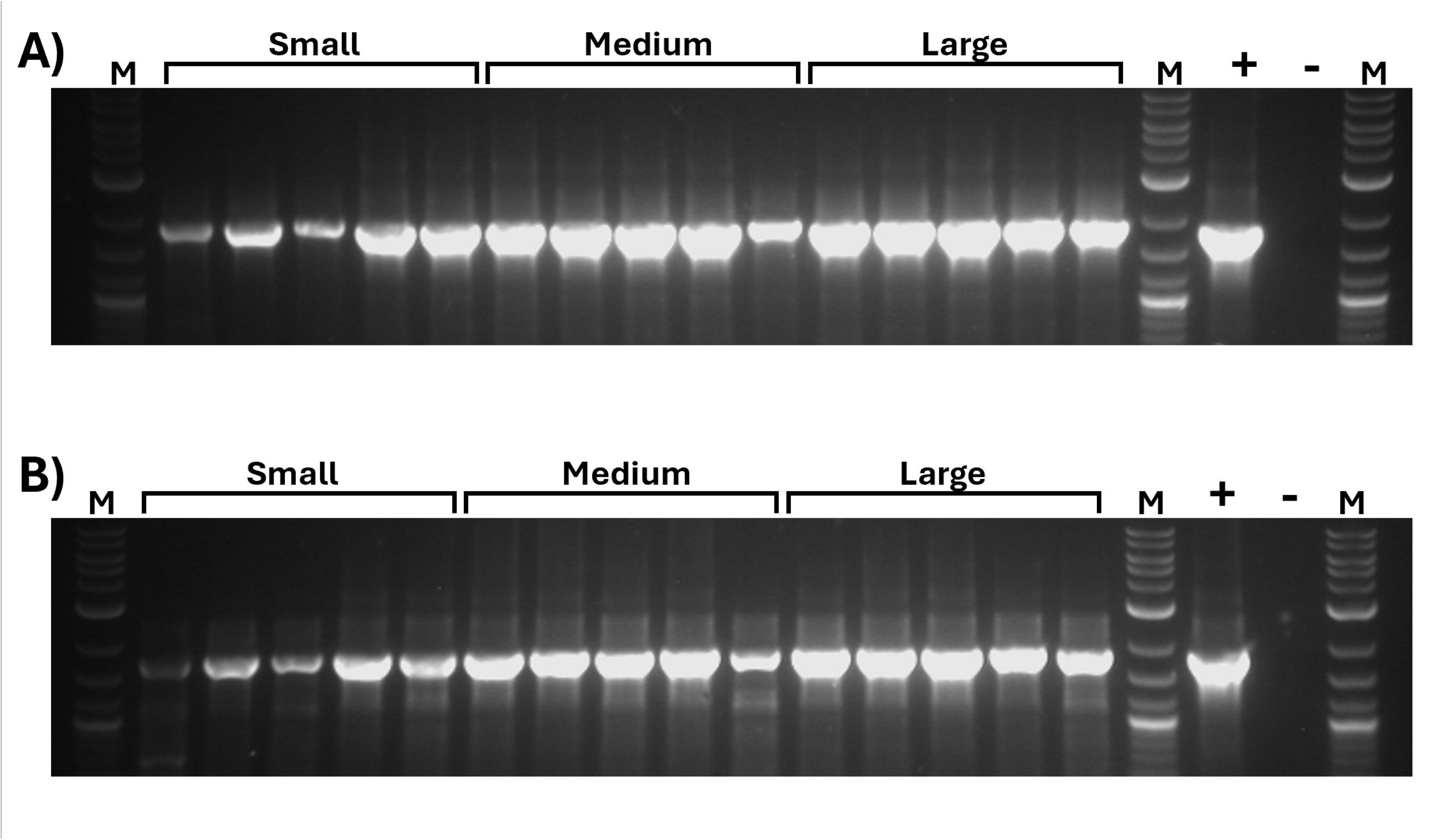

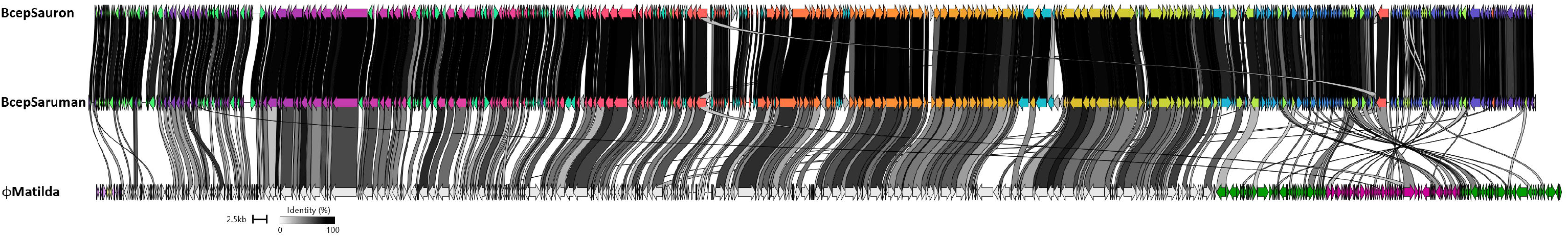

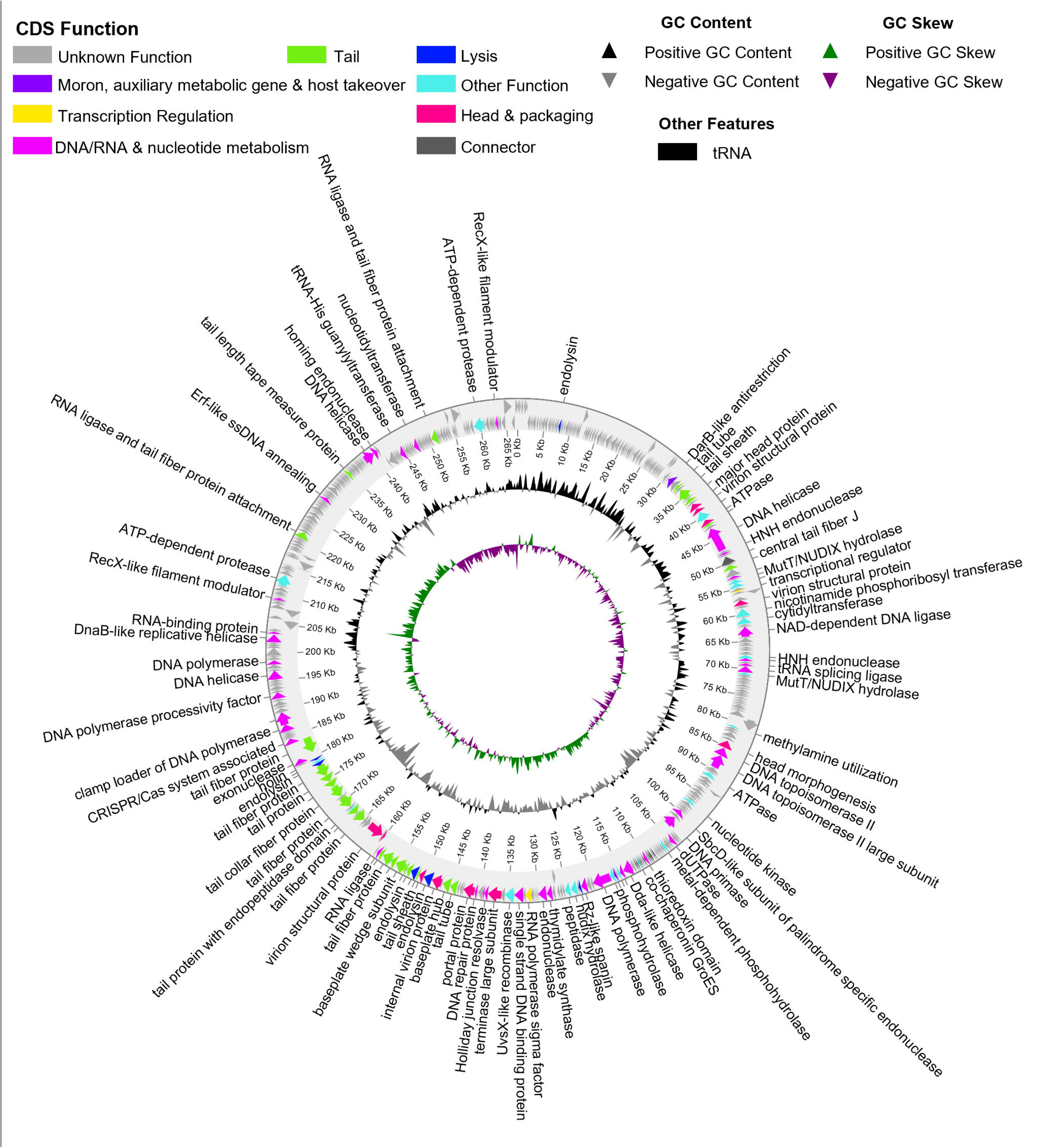

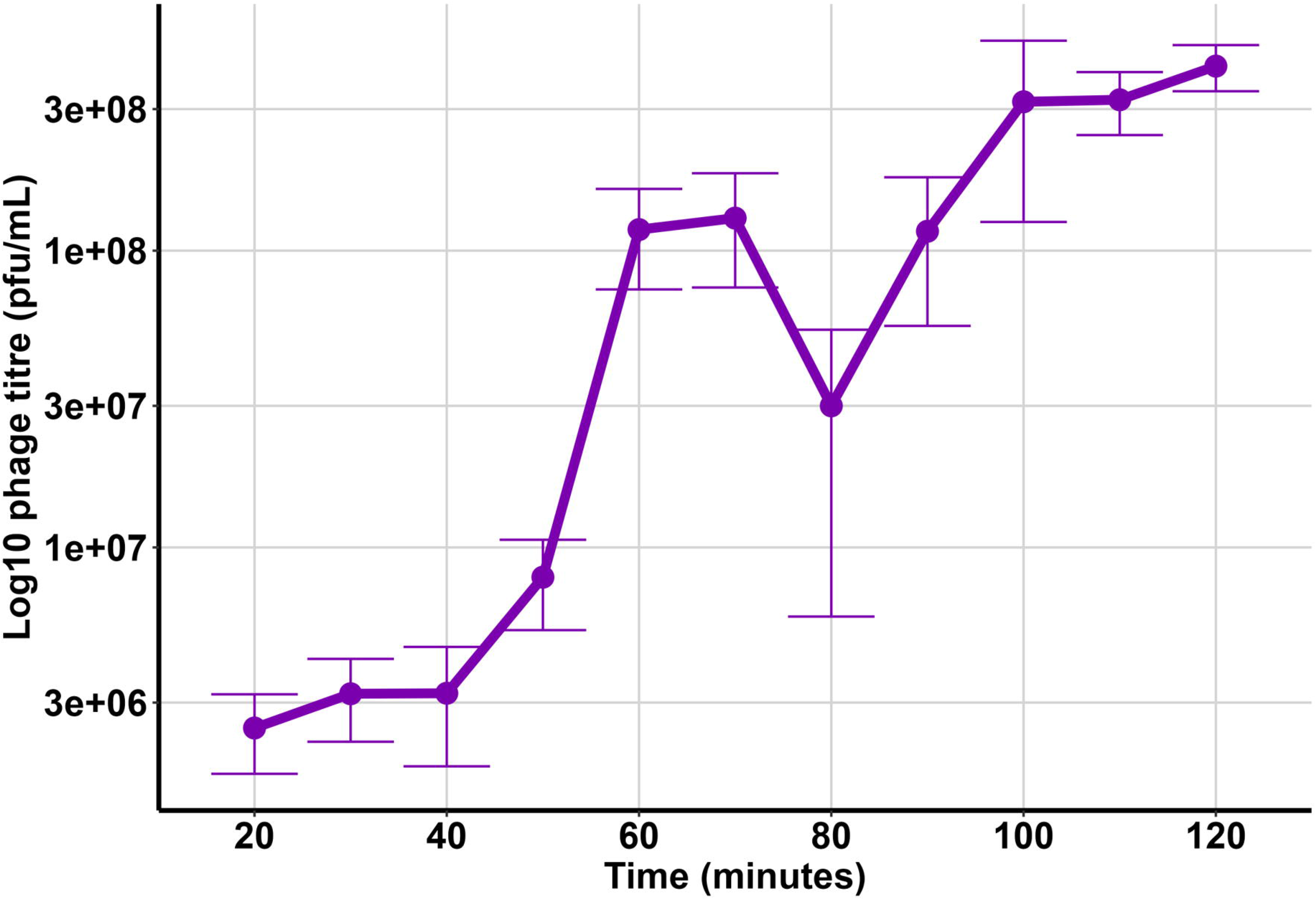

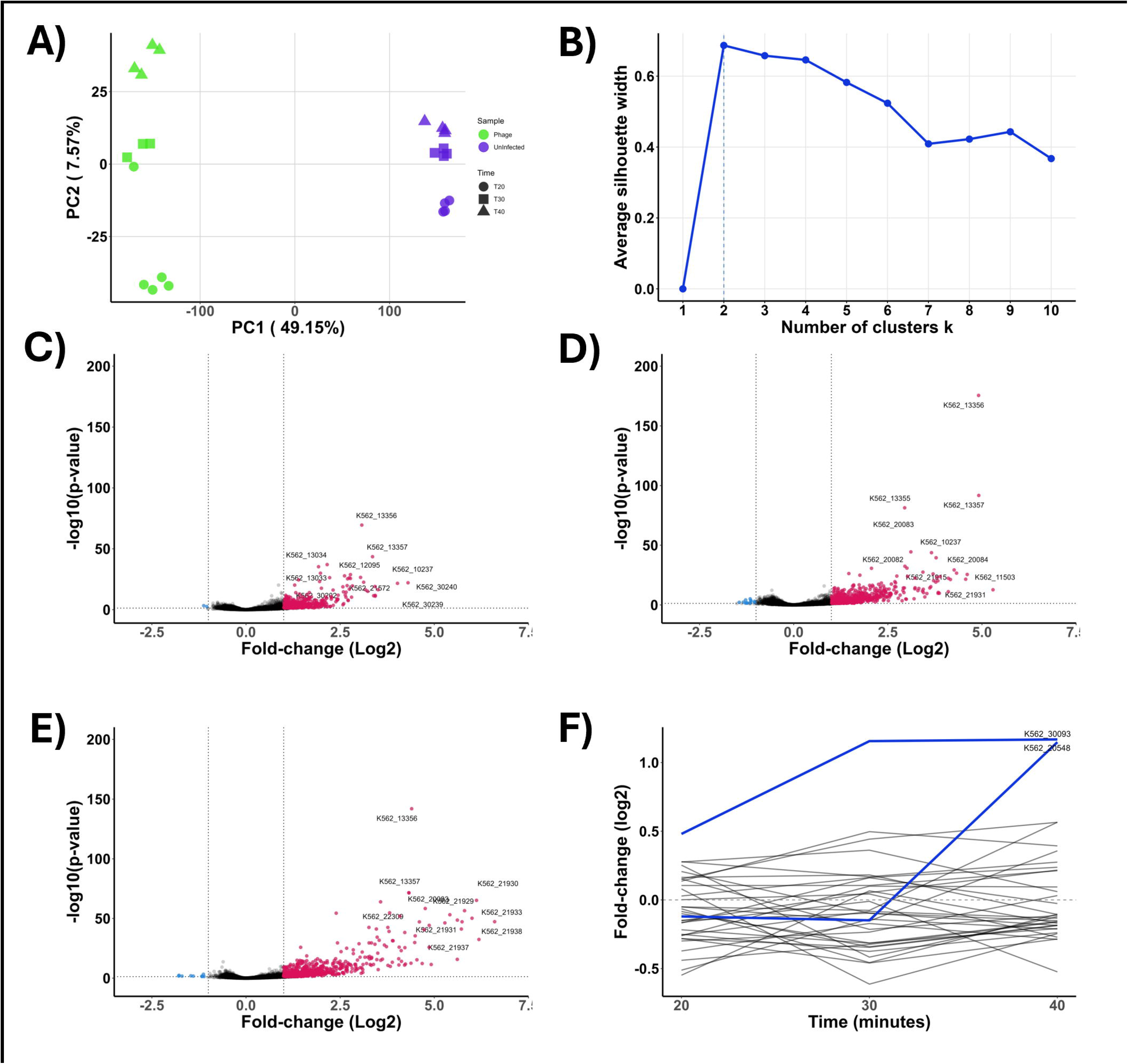

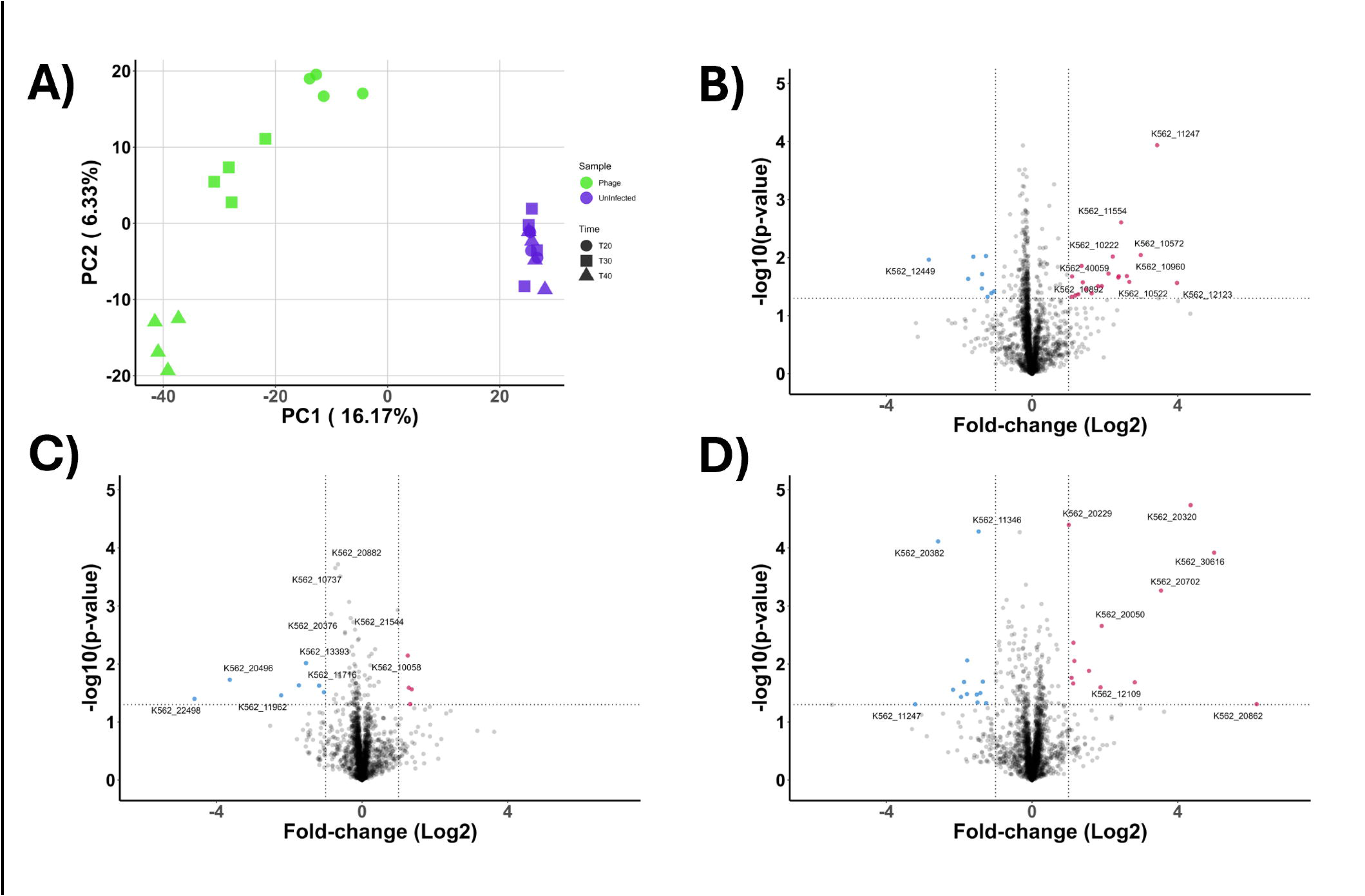

