## Supplementary material for "Unlocking the secrets of paired terminal genomic repeats utilised by the *Burkholderia* jumbo phage Matilda": Suplementary Data

#### Supplementary Tables

**Supplementary Table 1:** Bacterial strains used in this study. For BCC strains, the name in the brackets indicates the common strain name.

**Supplementary Table 2:** DNA primers used in this study.

**Supplementary Table 3:** Genome composition of φMatilda (attached)

**Supplementary Table 4:** DEseq2 comparisons of transcriptomics time course data containing the log2 fold change and standard error (lfcSE) and the native *p*-value and adjusted p-value (padj)

**Supplementary Table 5:** TPM counts of bacterial and phage genes

**Supplementary Table 6:** Proteomic time course data with label-free quantification intensity (LFQ.intensity) and paired comparison results.

### Supplementary Figures

#### Supplementary Figure 1

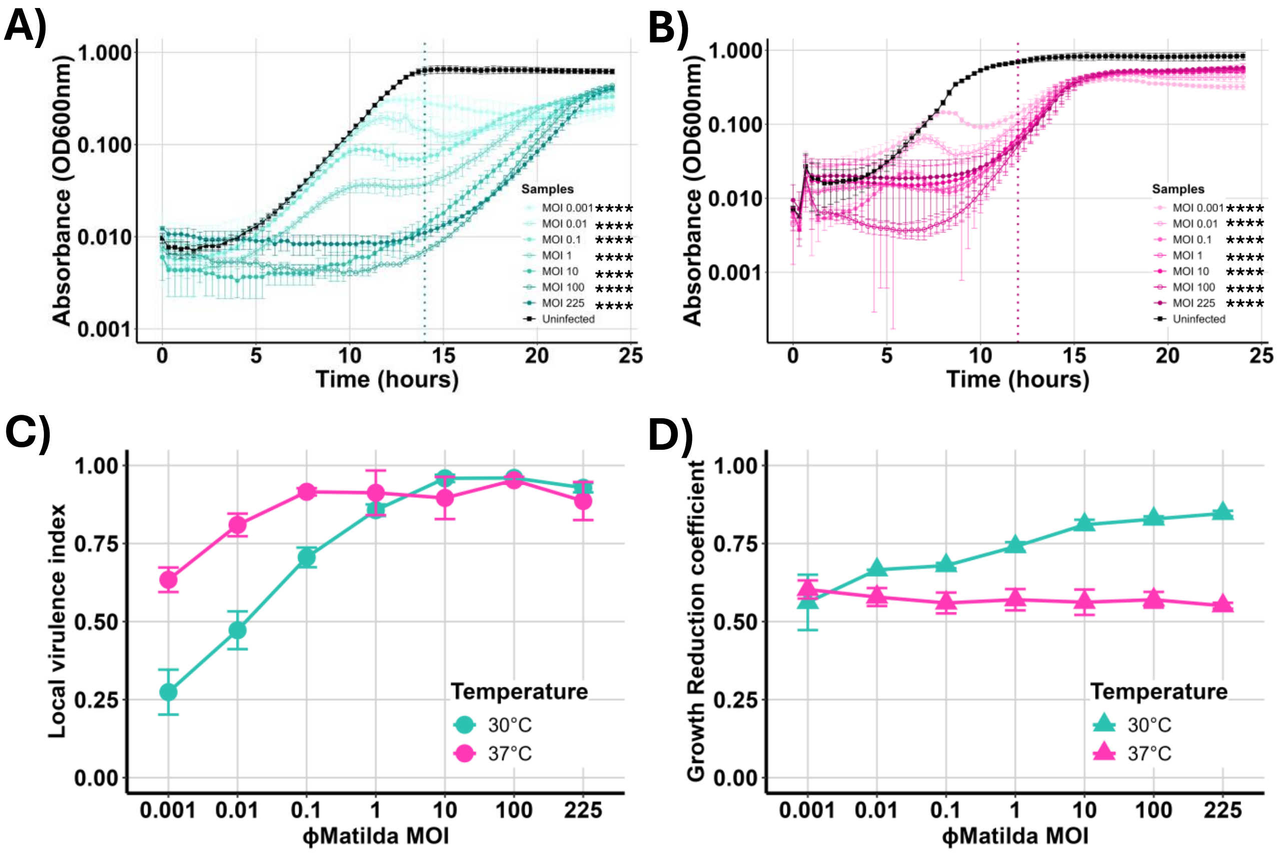


**Supplementary Figure 1: φMatilda killing kinetics.** Growth of *B. cenocepacia* K56-2 over 24 hours at **A)** 30°C and **B)** 37°C infected with varying concentrations of φMatilda. Statistical significance determined using a Kruskal-Wallis test is denoted next to the sample name. *p*-value <0.001 ****. The point at which stationary phase was reached is denoted with a dotted line. The area under the curve (AUC) before stationary phase was used to calculate the $V_i$ (**C**) relative to the uninfected sample. The AUC for the entire graph was used to calculate the GRC (**D**) relative to the uninfected sample. Data: mean ± SD, n=3.

**Supplementary Figure 2**

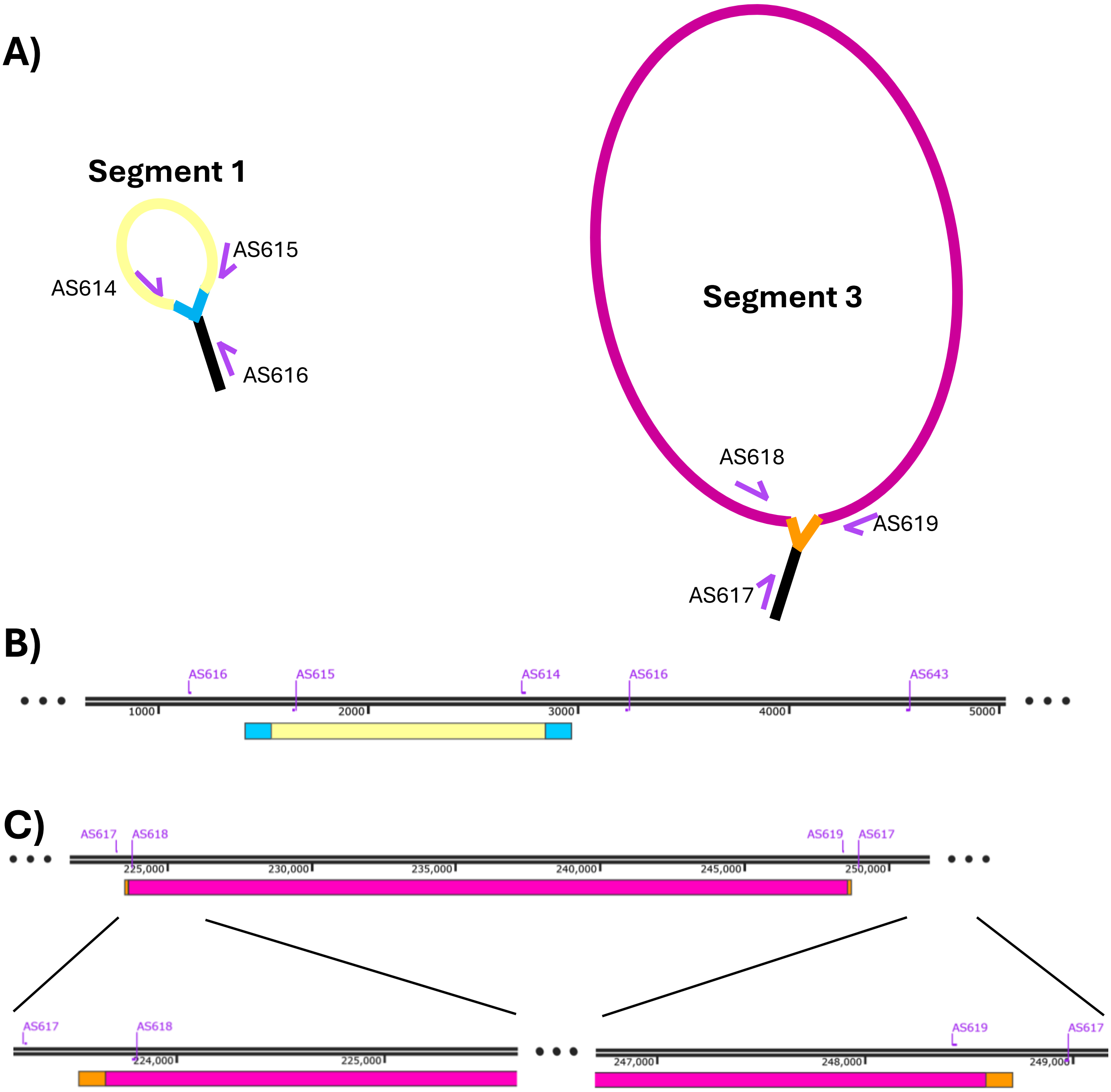


**Supplementary Figure 2: Schematic of primer binding sites to confirm joining of the three φMatilda genome segments**. **A)** Approximate binding sites of primers on Segment 1 (yellow), Segment 3 (pink), and the core genome (black) in the *de novo* assembly model. 127 bp repeats are annotated in blue for Segment 1 and orange for Segment 3. **B)** Linearised version of Segment *1 de novo* assembly model. **C)** Linearised version of Segment 3 *de novo* assembly model.

#### Supplementary Figure 3

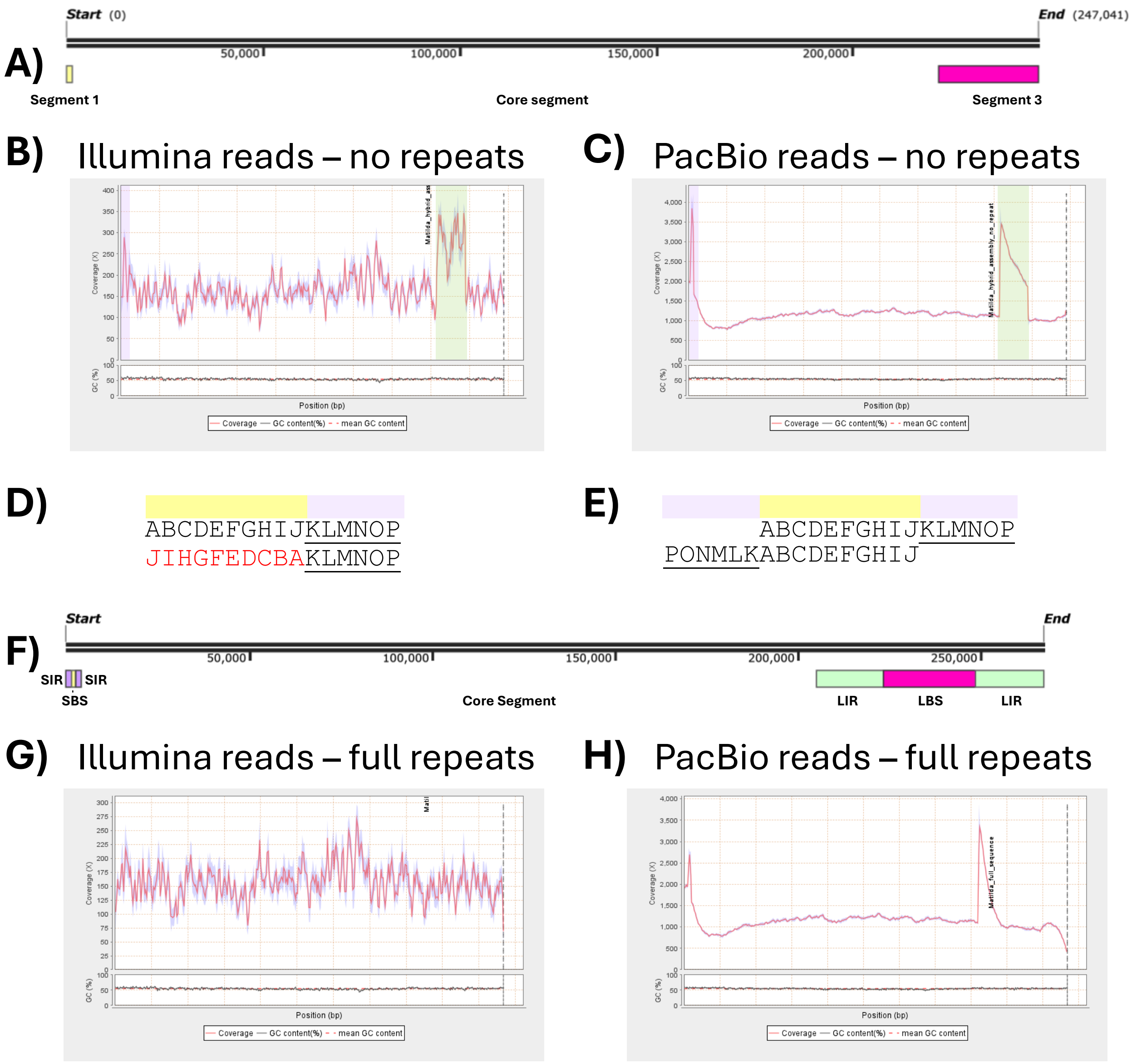


**Supplementary Figure 3: Genome schematic and sequencing read mapping of jumbo phage Matilda. A)** Linear map of φMatilda genome with the bidirectional segments presented in a fixed orientation. Total genome size 247,041 bp. Segment 1 is denoted with the yellow box and segment 3 is denoted with the pink box. **B)** Sequencing read coverage map of Illumina reads mapped against the 247,041 bp with a single orientation of the bidirectional sequences. Peaks due to repeated sequence shown in purple (short repeat) and green (long repeat). **C)** Sequencing read coverage map of PacBio reads mapped against the 247,041 bp with a single orientation of the bidirectional sequences. Peaks due to repeated sequence shown in purple (short repeat) and green (long repeat). **D)** Example of partially mapped read sequence. The sequence in black is correctly mapped. The sequence in red is incorrectly mapped. Underlining/purple represents where the peak is occurring. **E)** Example of the sequences correctly mapped. Underlining/purple represents the inverted repeat sequence. **F)** Linear map of φMatilda genome with the bidirectional segments presented in a fixed orientation. Total genome size 266,947 bp. The SBS is denoted with the yellow box and LBS is denoted with the pink box. The short and long inverted repeats are denoted with the purple and pink boxes respectively. **G)** Sequencing read coverage map of Illumina reads mapped against the 266,947 bp with a single orientation of the bidirectional sequences. **H)** Sequencing read coverage map of PacBio reads mapped against the 266,947 bp with a single orientation of the bidirectional sequences. Peaks due to repeated sequence shown in purple (short repeat) and pink (long repeat).

**Supplementary Figure 4**

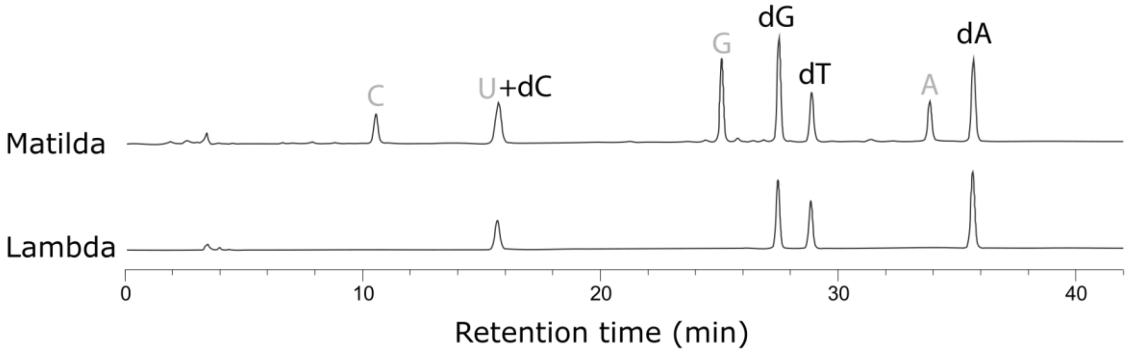


**Supplementary Figure 4**: **LC-MS chromatogram of digested φMatilda and λ phage DNA.** Peaks corresponding to either deoxyribonucleotides (black) or ribonucleotides (grey) are annotated.

**Supplementary Figure 5**

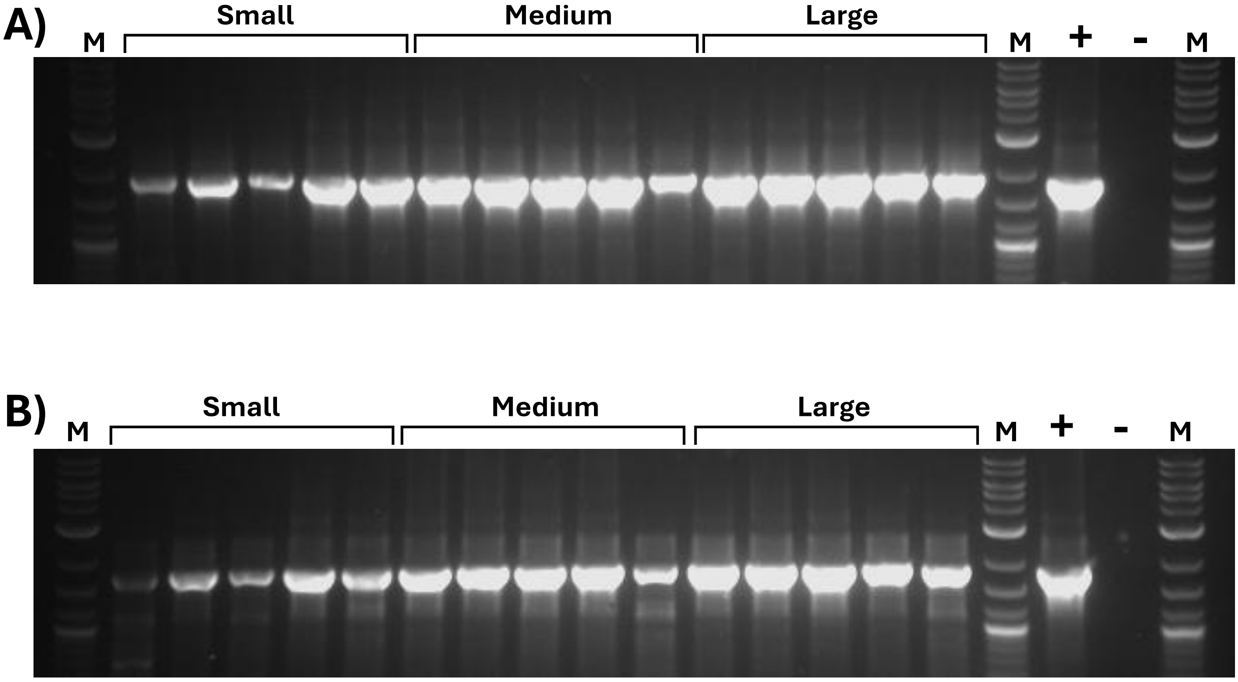


**Supplementary Figure 5: Confirmation of the presence of the small bidirectional segment being attached to the core genome in both orientations in various plaque sizes.** Five small (<0.5 mm), medium (0.5-1 mm), and large (>1 mm) φMatilda plaques were screened for the orientation of the small bidirectional segment using **A)** AS614/AS643 (Orientation 1) and **B)** AS615/AS643 (Orientation 2). The 1 kb plus NEB ladder is denoted with M. Purified φMatilda DNA was used as a positive control (+) and a No DNA control (-) was also included.

#### Supplementary Figure 6

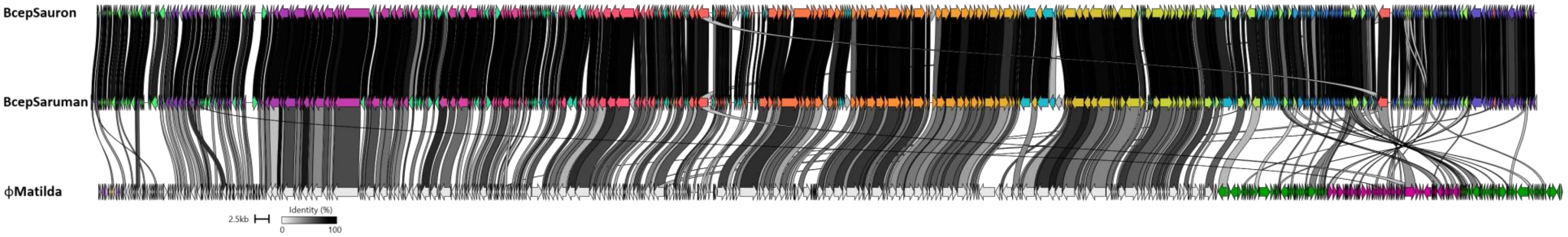


**Supplementary Figure 6: Genomic comparison of φMatilda and the Sarumanviruses BcepSauron and BcepSaruman.** The coding sequences (CDSs) of the Sarumanviruses are colour-coded based on shared homology. The CDSs of φMatilda are coloured to reflect the genome architecture: Purple – short inverted repeats, yellow – small bidirectional segment, grey – core genome, green – long inverted repeat, and pink – large bidirectional segment. The links between the phages are coloured to reflect their shared identity (legend on bottom left). Scale bar represents 2.5 kb.

#### Supplementary Figure 7

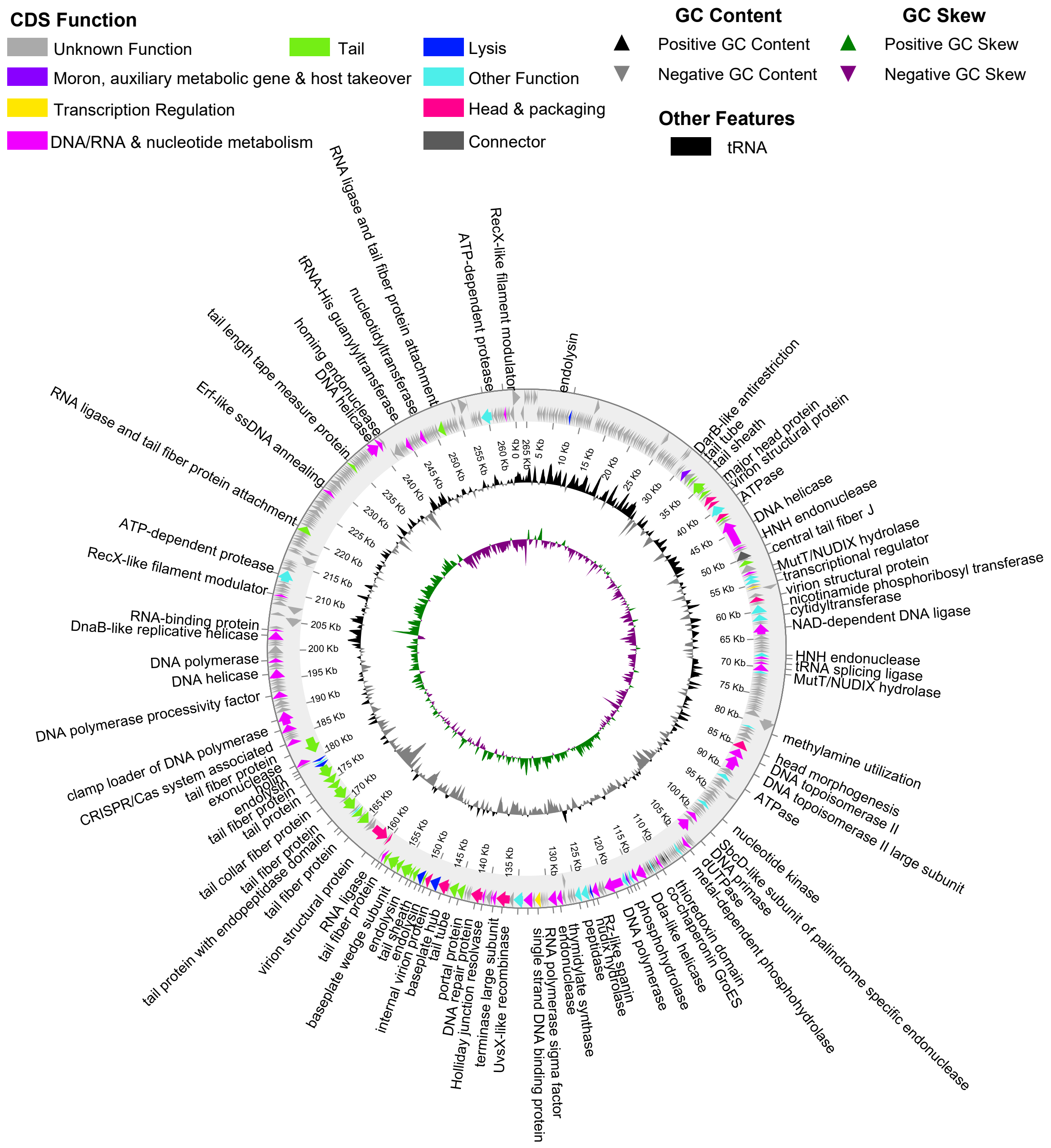


**Supplementary Figure 7: Graphical representation of the φMatilda genome.** The outer ring shows the CDSs with the colour corresponding to the predicted function (legend above). tRNAs are annotated in black. The middle ring shows the GC content: black - positive GC content, grey - negative GC content. The inner ring denotes the GC skew: purple - positive GC skew, green - negative GC skew.

Supplementary Figure 8

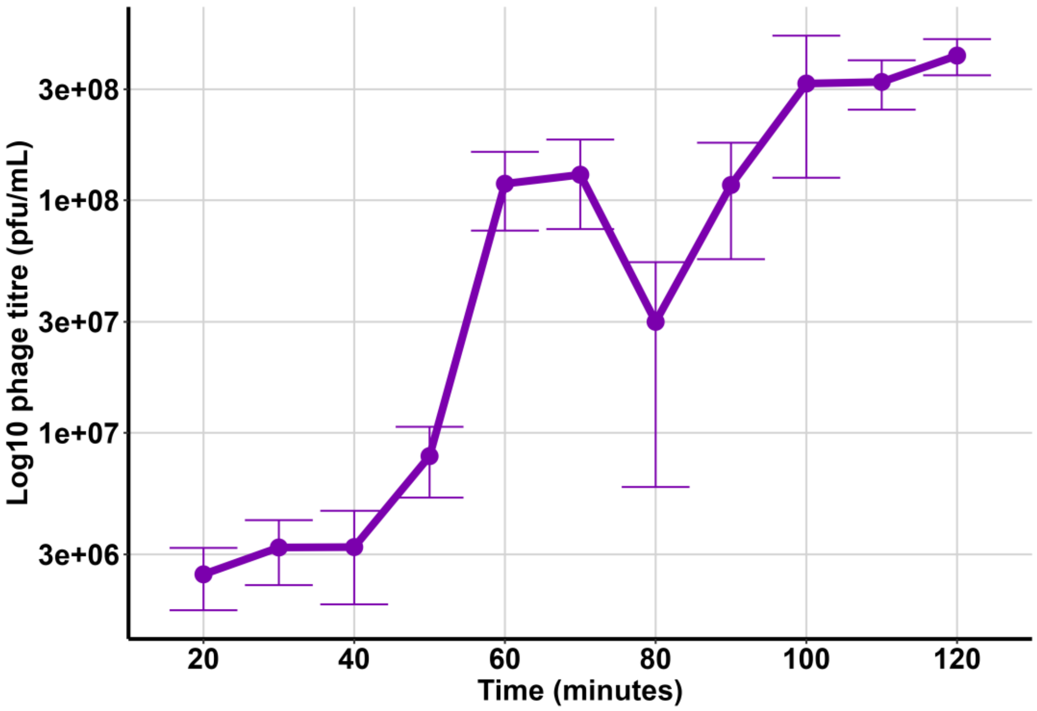


**Supplementary Figure 8: One-step growth curve of φMatilda using wildtype** *B. cenocepacia* K56-2 at an MOI of 3. The mean ± SD of the phage titre at each time point over 100 minutes is shown. N=4.

#### Supplementary Figure 9

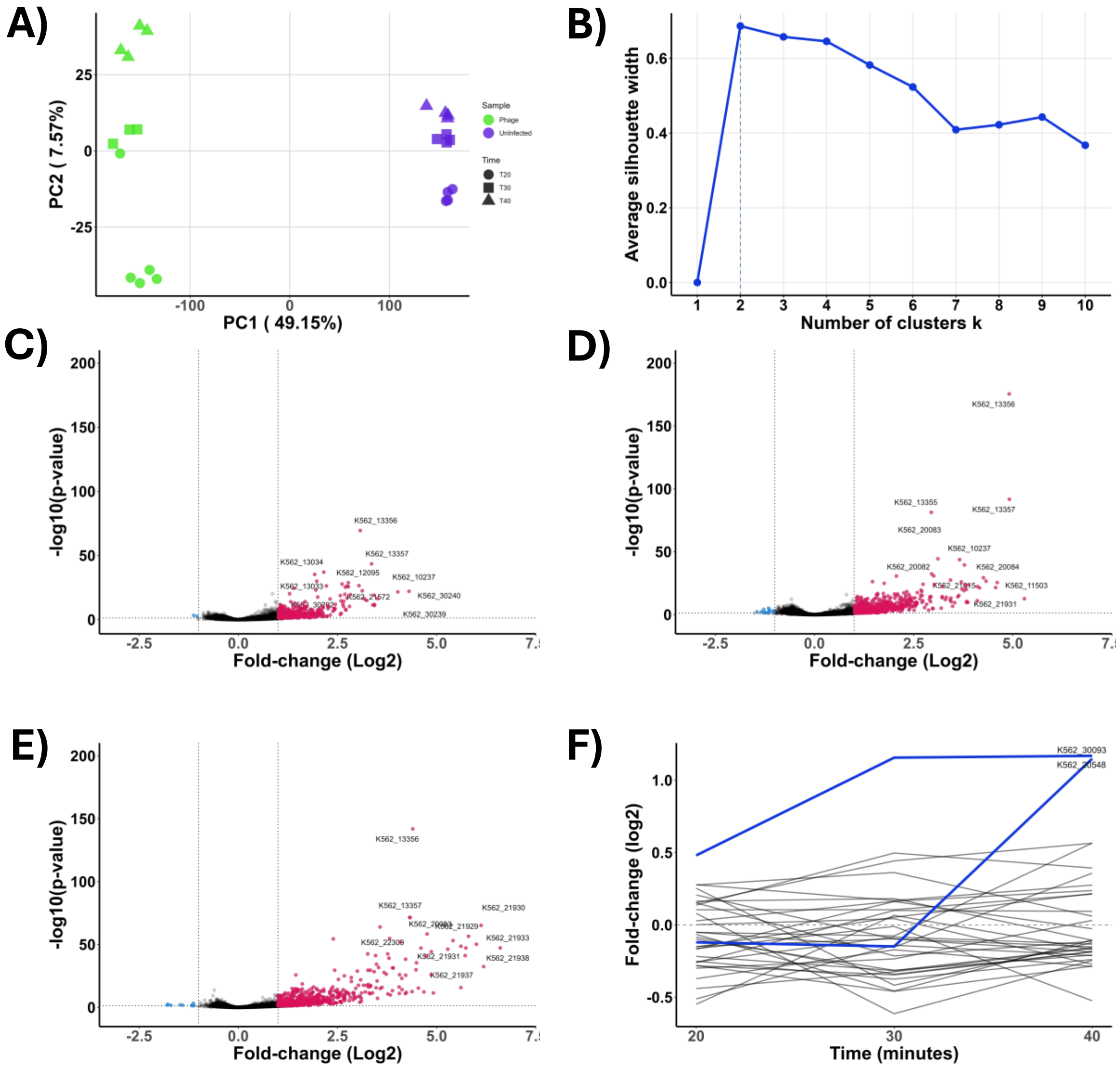


**Supplementary Figure 9: Transcriptomic validation and host response. A)** PCA revealed clear separation between infected and uninfected samples. **B)** Silhouette analysis of φMatilda gene expression across infected samples used to determine the optimal number of transcriptional clusters. **C-E)** Volcano plot showing differential expression of *B. cenocepacia* K56-2 genes between uninfected and infected samples at time 0 (**C**), time 10 (**D**), and time 20 (**E**). The X-axis shows the $log_2$ fold-change and the y-axis shows the $-log_{10}$(adjusted *p*-value). Genes with significantly increased expression are shown in pink with the top 10 most significantly differentially expressed genes are labelled. Genes with significantly decreased expression in the infected samples are shown in blue. **F)** Line plot of the genes associated with the anti-phage defence systems in *B. cenocepacia* K56-2. The two genes showing significant differential expression at 20 minutes are highlighted in blue.

**Supplementary Figure 10**

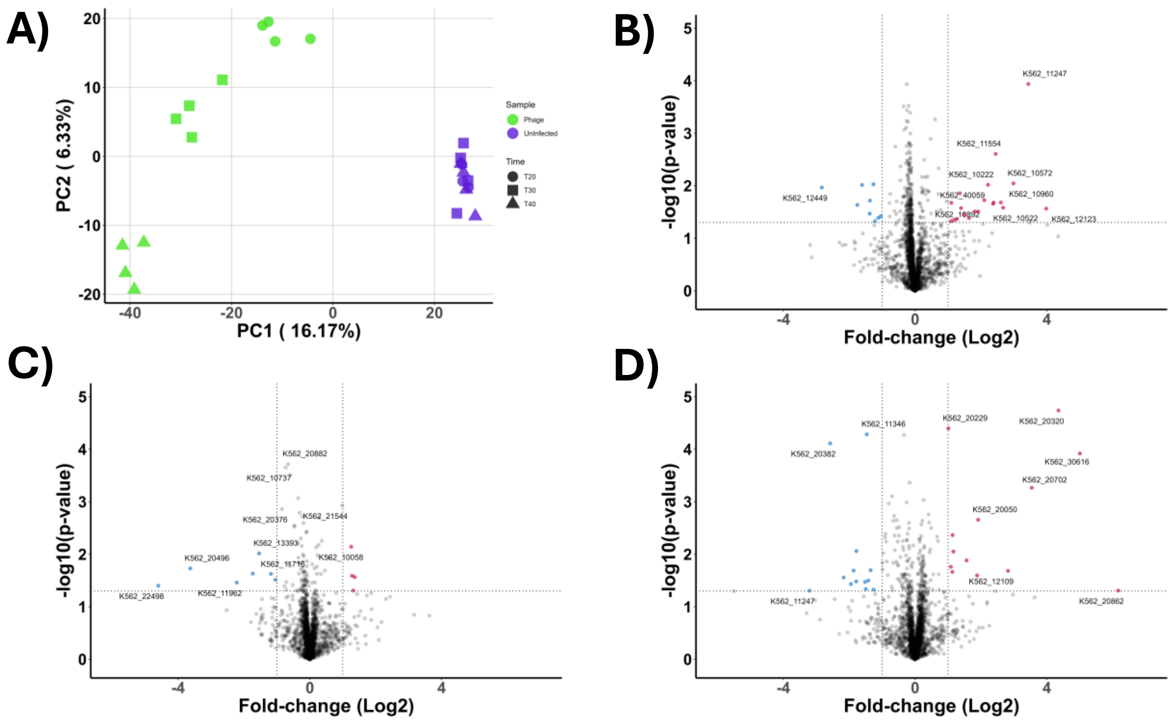


**Supplementary Figure 10: Proteomic validation and host response. A)** PCA revealed clear separation between infected and uninfected samples. **B-D)** Volcano plot showing differential production of *B. cenocepacia* K56-2 proteins between uninfected and infected samples at time 0 (**B**), time 10 (**C**), and time 20 (**D**). The X-axis shows the $\log_2$ fold-change and the y-axis shows the $-\log_{10}$(*p*-value). Proteins with significantly increased production are shown in pink. Proteins with significantly decreased production in the infected samples are shown in blue. The top 10 most significantly differentially produced proteins labelled.
